# The Temporal Dynamics of Aperiodic Neural Activity Track Changes in Sleep Architecture

**DOI:** 10.1101/2024.01.25.577204

**Authors:** Mohamed S Ameen, Joshua Jacobs, Manuel Schabus, Kerstin Hoedlmoser, Thomas Donoghue

## Abstract

The aperiodic (1/f-like) component of electrophysiological data - whereby power systematically decreases with increasing frequency, as quantified by the aperiodic exponent - has been shown to differentiate sleep stages. Earlier work, however, has typically focused on measuring the aperiodic exponent across a narrow frequency range. In this work, we sought to further investigate aperiodic activity during sleep by extending these analyses across broader frequency ranges and considering alternate model definitions. This included measuring ‘knees’ in the aperiodic component, which reflect bends in the power spectrum, indicating a change in the exponent. We also sought to evaluate the temporal dynamics of aperiodic activity during sleep. To do so, we analyzed data from two sources: intracranial EEG (iEEG) from 106 epilepsy patients and high-density EEG from 17 healthy individuals, and measured aperiodic activity, explicitly comparing different frequency ranges and model forms. In doing so, we find that fitting broadband aperiodic models and incorporating a ‘knee’ feature effectively captures sleep-stage-dependent differences in aperiodic activity as well as temporal dynamics that relate to sleep stage transitions and responses to external stimuli. In particular, the knee parameter shows stage-specific variation, suggesting an interpretation of varying timescales across sleep stages. These results demonstrate that examining broader frequency ranges with the more complex aperiodic models reveals novel insights and interpretations for understanding aperiodic neural activity during sleep.

## 1. Introduction

Studies investigating the electrophysiology of sleep are classically based on a scoring system focusing on oscillatory activity (such as alpha oscillations, sleep spindles and slow wave activity) to categorize sleep into different stages (Berry et al., 2017). The electrophysiological signal, however, contains a mixture of oscillatory, periodic components that rise above a non-oscillatory, aperiodic signal (Freeman & Zhai, 2009; He et al., 2010; Miller et al., 2009). The aperiodic signal decays with increasing frequency in a 1/f^x^ relationship, quantifiable by the aperiodic exponent (x), which corresponds to the slope on the log-log power spectrum (He et al., 2010; Miller et al., 2009; Voytek et al., 2015). Building on earlier research that identified differences in aperiodic activity between sleep and wake states (Feinberg et al., 1984; Freeman & Zhai, 2009; He et al., 2010; Pereda et al., 1998), recent studies have consistently found that the spectral exponent varies across different sleep stages (G. Horváth et al., 2022; Höhn et al., 2023; Kozhemiako et al., 2022; Lendner et al., 2020, 2023; Miskovic et al., 2019; Rosenblum et al., 2022; Schneider et al., 2022), contributing to a growing body of literature focusing on aperiodic brain activity during sleep (Figure 1A).

**Figure 1.**
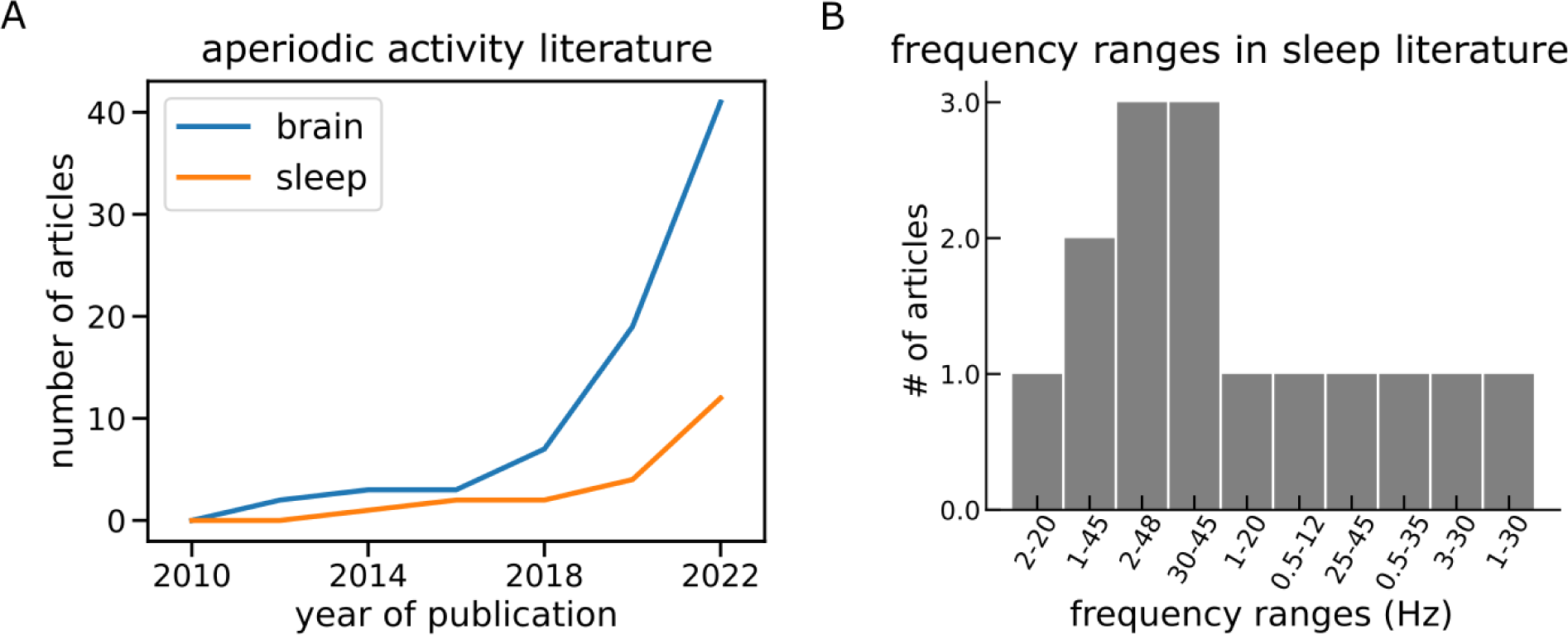
Review of literature examining aperiodic activity during sleep. A) The number of peer-reviewed articles investigating aperiodic activity in the brain and during sleep from 2010 till 2022. B) A summary of the frequency ranges used to measure aperiodic brain activity in the sleep literature.

The observed changes in aperiodic activity during sleep provide potential information about the underlying physiological changes. Computational models, supported by empirical data from animal studies, have proposed the spectral exponent as a non-invasive measure of cortical excitation-Inhibition (E-I) ratio (Chini et al., 2022; Gao et al., 2017; Lendner et al., 2023; Lombardi et al., 2017; Trakoshis et al., 2020), with a steeper exponent indicating stronger inhibition and a flatter exponent reflecting more excitation. From wakefulness to non-rapid eye movement (NREM) sleep the exponent of the electroencephalography (EEG) signal becomes steeper, mirroring the increase in inhibition reported during sleep (Niethard et al., 2016; Birdi et al., 2020), and subsequently becomes slightly flatter during rapid eye movement (REM) sleep (Lendner et al., 2020; Hoehn et al., 2022, Kozhemiako et al., 2022; Rosenblum et al., 2022).

Accurately measuring and interpreting aperiodic activity requires methods that effectively differentiate aperiodic and oscillatory activity, which need to be employed using appropriate settings (Donoghue & Watrous, 2023; Gerster et al., 2022). A key decision when examining aperiodic neural activity includes selecting the frequency range to analyze. Previous sleep studies have thus far examined a wide variety of frequency ranges (Figure 1B), with a tendency towards examining a narrow frequency band within the range of 30 to 45 Hz (Demirel et al., 2021; Höhn et al., 2023; Kozhemiako et al., 2022; Lendner et al., 2020), a range chosen due to its potential relationship to the E-I ratio (Gao et al., 2017). In addition to the frequency range, analyzing aperiodic neural activity requires choosing the appropriate model form by selecting the function to fit to the power spectrum (Figure 2). The vast majority of the existing literature in sleep research has applied measures of a single exponent fit, such that overall existing sleep studies have tended to examine a simple model fit to a narrow range, only characterizing and describing a restricted view of the neural power spectrum.

**Figure 2.**
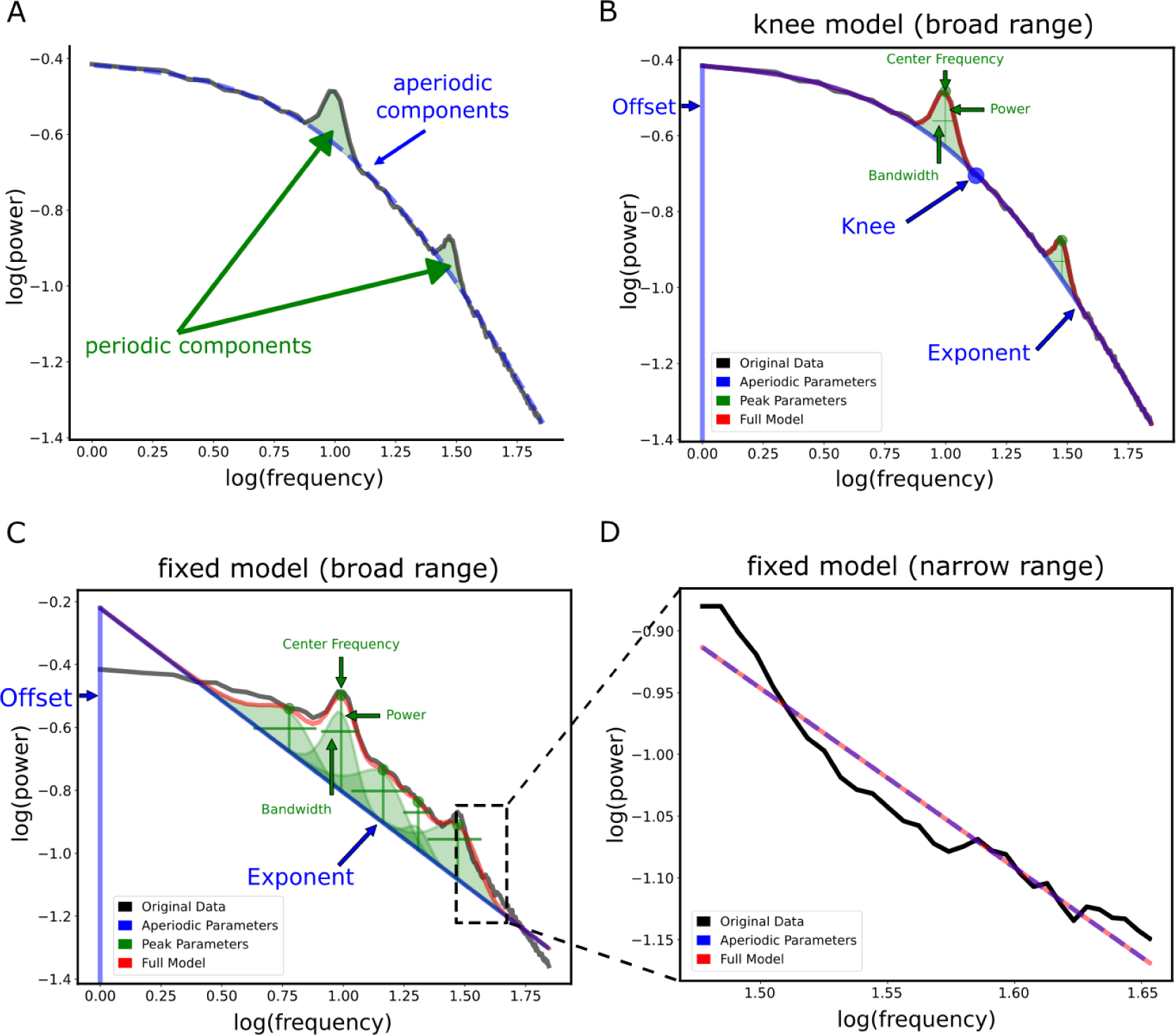
Schematic of measuring aperiodic activity from power spectra, examining different frequency ranges and model forms. A) An example simulated power spectrum, with 2 oscillatory peaks at 10 Hz and 30 Hz, a knee at 13.13 Hz and an exponent of 1.25. Periodic components are highlighted in green, and the simulated aperiodic component is shown in the dashed blue line. B) An example spectral model fit using a ‘knee’ model, i.e. a model that incorporates a knee parameter, with annotated spectral features. Notably, fitting this model over a broad frequency range (1-45 Hz) produced very high Goodness of fit measure (R^2^ = 0.99) and low mean squared error (MSE = 0.004). C) Example spectral model fit using the same frequency band but fitting a fixed model, i.e. a model that assumes a fixed exponent value, resulted in a high R^2^ (0.99) but increased error (MSE = 0.013). Note the difference in the number of oscillatory peaks (green) between B and C - while both models can attain high R^2^, if there is a model mismatch, the fixed model tends to overfit oscillatory peaks. D) An example model fit of a fixed model fit over a narrow (30-45 Hz) frequency band. This model had the lowest model performance (R^2^ = 0.95, MSE = 0.016). For an example of fitting a model without simulated a knee component see Suppl. Figure 2-1, which shows that again, both the knee and fixed models showed high and comparable performance levels (both with an R^2^ of 0.99), in contrast to the narrowband model fit which had a lower fit value (R^2^ = 0.95).

A key potential strength of analyzing aperiodic activity is to be able to characterize broad frequency ranges. To do so, one has to assess the overall shape of the power spectral density (PSD) to choose the most appropriate frequency range and model form. Notably, the neural power spectrum often displays an inflection point, referred to as the “knee” (Miller et al., 2009; Gao et al., 2020). The frequency at which this knee occurs, i.e. the knee frequency, has been proposed to reflect the population timescale (Miller et al., 2009; Gao et al., 2020) and has also been shown to change across sleep stages (Lender et al., 2023). Collectively, this suggests that analyses of aperiodic neural activity during sleep may benefit from formalizing and potentially extending the frequency range under study. To do so, however, requires considering different fit functions (e.g., fitting a function with a knee parameter), and choosing appropriate frequency ranges to fit across.

In addition, most studies of aperiodic activity during sleep thus far have predominantly focused on the examination of aperiodic activity over entire sleep stages. Recent methodological and empirical developments have begun to demonstrate the rich temporal dynamics of aperiodic activity and their relation to behavior (Donoghue et al., 2020; Waschke et al., 2021; Wilson et al., 2022). This suggests that previous analyses may have overlooked nuanced dynamics within and across sleep stages. However, novel methods are now available that can now be used to investigate questions such as whether changes in aperiodic activity shows sharp transitions or slow drifts between sleep stages, and explore whether aperiodic activity undergoes event-related changes during sleep.

Based on these considerations, our objective was to expand upon previous research by broadening the investigation of aperiodic activity during sleep. Specifically, our approach involved explicitly evaluating different frequency ranges and model forms in order to select measurements of aperiodic activity that accounts for the full shape of the PSD and captures nuances like the knee frequency. We then applied these measures in a time-resolved manner, in order to examine sleep dynamics across multiple temporal scales during the night. To do so, we analyzed publicly available intracranial EEG (iEEG) data from across different sleep stages (von Ellenrieder et al, 2020) as well as a high-density EEG dataset from healthy human participants through an entire night of sleep (Blume et al., 2018). To measure aperiodic neural activity, we used the specparam toolbox-formerly ‘FOOOF’- (Donoghue et al., 2020) to compare model forms, fit spectral models, and compute time-resolved estimates.

In doing so, we highlight the advantages of employing a model form for aperiodic activity that examines broader frequency ranges and accounts for the presence of the knee, which we find to be a prominent feature in sleep recordings. Our findings demonstrate that the knee frequency systematically relates to different sleep stages, and suggest a connection between the changes in knee frequency across different sleep stages observed here and the corresponding variations in the exponent values highlighted by previous work. Furthermore, by applying this model in a temporally resolved manner, we demonstrate that the temporal dynamics of aperiodic activity track transitions between sleep stages, and also respond to external stimuli. Collectively, this work demonstrates that by expanding the model forms and temporal resolution of aperiodic activity measures, we can capture and characterize more of the variance of sleep data, in a way that both highlights systematic relationships to sleep architecture and offers putative interpretations of the underlying neural circuits.

## 2. Methods

### 2.1. Literature search

To systematically examine the prior literature on aperiodic activity during sleep, we used the open source toolbox LISC (v0.3.0) in python (Donoghue, 2019) to collect and analyze scientific articles. We measured the co-occurrences of the term: ‘brain’, or the term: ‘sleep’ with terms that indicate aperiodic activity: ‘spectral slope’, ‘spectral exponent’, ‘1/f exponent’, ‘1/f slope’, ‘aperiodic slope’, ‘aperiodic exponent’, or ‘power-law exponent’. We also added inclusion terms to limit the search to only electrophysiological studies by adding the terms: ‘EEG’, ‘MEG’, ‘iEEG’, ‘intracranial EEG’, ‘electroencephalography’, ‘magnetoencephalography’, and ‘intracranial electroencephalography’. We collected the number of articles containing these terms in their titles or abstracts from the Pubmed database, for the time range of 2009-2023, in 2 year bins. To collect the frequency ranges previously analyzed in the sleep literature, we used the same terms to search for studies examining aperiodic activity with the inclusion term ‘sleep’. We included articles that a) investigated aperiodic activity in humans, and b) had a clear mention of the frequency range used to fit the model. The set of papers that matched these criteria included 15 articles (Table 1). Subsequently, we added three early publications investigating aperiodic activity during sleep which were not detected by the algorithm due to using a different label for the exponent. We manually examined these articles to identify the specific frequency ranges used for estimating aperiodic activity.

**Table 1.** The results of the literature search. A list of publications that investigated the spectral exponent in the human brain during sleep. These articles had a clear mention of the frequency range used to fit the model. Other studies which did not explicitly mention the range were not included. A single study (Maschke et al., 2023), incorporated the use of a knee model in its analysis. This research reported an enhanced model fit when utilizing the knee model compared to the fixed model.

| Article | Frequency band |
| --- | --- |
| Alnes et al., 2023 | 2 - 20 Hz |
| Ameen et al., 2023 | 1 - 45 Hz |
| Bódizs et al., 2021 | 2 - 48 Hz |
| Demirel et al., 2021 | 30 - 45 Hz |
| Favaro et al., 2023 | 1 - 20 Hz |
| Feinberg et al., 1984 | 0.5 - 12 Hz |
| G. Horváth et al., 2022 | 2 - 48 Hz |
| Kozhemiako et al., 2022 | 30 - 45 Hz |
| Lendner et al., 2020 | 30 - 45 Hz |
| Lendner et al., 2023 | 25 - 45 Hz |
| Maschke et al., 2023 | 1 - 45 Hz |
| Miskovic et al., 2019 | 0.5 - 35 Hz |
| Pereda et al., 1998 | 3 - 30 Hz |
| Schneider et al., 2022 | 2 - 48 Hz |
| Wen and Liu, 2016 | 1 - 30 Hz |

### 2.2. Datasets

#### 2.2.1. iEEG data

We analyzed an openly accessible and previously published dataset from the Montreal Neurological Institute (MNI; https://mni-open-ieegatlas.research.mcgill.ca/). The dataset contains one minute recordings from 106 patients (52 females, 33.1 ± 10.8 years) with focal epilepsy during quiet wakefulness with eyes closed, NREM sleep (N2 and N3), and REM sleep. The dataset contains 1772 channels during Wake (Frauscher et al., 2018), 1468 channels in NREM sleep and 1012 channels in REM sleep (von Ellenrieder et al., 2020).

##### Preprocessing

Detailed preprocessing steps are described in the associated paper (von Ellenrieder et al., 2020). Briefly, raw data were low-pass filtered at 80 Hz, resampled to 200 Hz, and had line noise removed. Thereafter, the data were visually inspected by an expert neurophysiologist and artifactual segments of the signal were removed. Consequently, for some patients, the data used to compile one minute of recording comprised several non-consecutive segments. Each channel in each segment was demeaned. When segments were concatenated, a buffer time of 2 s of zeros was added between the two segments. All the channels were then zero padded at the end to a length of 68 s if necessary to have a uniform length regardless of the number of segments. The dataset contains signals organized into 38 brain regions, with each region containing data from five patients. The five patients differed between regions but were always the same across sleep stages per region. For our analysis, we excluded 4 non-cortical regions, namely: the hippocampus, the amygdala, the fusiform and parahippocampal gyri. Thus, we report the results from 34 cortical regions.

#### 2.2.2. EEG data

We also analyzed a dataset of high density EEG from 17 healthy human participants (14 females, 22.6 years ± 2.3) who spent two nights with high-density EEG in the sleep laboratory at the University of Salzburg (Blume et al., 2018). The first night served as an adaptation night, in which we recorded polysomnography (PSG) data. The second night was an experimental night, during which we recorded PSG data while also presenting sounds throughout the night. Here we used data only from the experimental night. The sounds were the subject’s own name (SON) and two unfamiliar names (UNs) spoken by either a familiar voice (FV) to the subject or an unfamiliar voice (UFV). The stimuli were played via loudspeakers in the room and started immediately when the subjects went to bed. Throughout the night, auditory stimuli were presented continuously for 90 minutes (stimulation periods) then followed by a 30-minute phase of quiet sleep (no-stimulation periods). This accounts for a 120-minute cycle that was repeated four times during the night. We adjusted the sound volume individually for each participant based on their preference, ensuring that it was clearly audible without being excessively loud to avoid disrupting their sleep. We jittered the inter-stimulus interval between 2800 and 7800 ms in 500 ms steps. At the beginning of the experiment, all participants signed written informed consents. The ethics department of the University of Salzburg approved the experiment.

##### Acquisition and preprocessing

We used a high-density EEG 256-channel GSN HydroCel Geodesic Sensor Net (Electrical 478 Geodesics Inc., Eugene, Oregon, USA) and a Net Amps 400 amplifier. Data were acquired at a sampling rate of 250 Hz with Cz as the online reference. For preprocessing, face and neck channels (53 channels) were excluded and the analyses were carried out on 183 EEG channels. We high-pass-filtered the signal from these channels at 0.1Hz, then we removed the 50 Hz line noise using a notch filter. Using the PREP pipeline (Bigdely-Shamlo et al., 2015), we removed and interpolated bad channels, restored the reference electrode (Cz), and re-referenced the data to an average reference. Finally, we performed independent component analysis (ICA) in EEGLab and visually labeled and removed noise-, heart-, and eye-related components.

#### 2.2.3. Simulated data

For example demonstrations of fitting spectral models, we simulated an example neural power spectrum using the specparam toolbox (Donoghue et al., 2020) with a frequency range of 1-45 Hz including two oscillatory peaks at 10 and 30 Hz, a knee frequency at 13.13 Hz, an exponent value of 1.25.

### 2.3. Sleep staging and K-complex (KC) detection

Sleep EEG data were scored semi-automatically using an algorithm developed by the Siesta group (Somnolyzer 24×7; The SIESTA Group Schlafanalyse GmbH., Vienna, Austria; Anderer et al., 2005; Anderer et al., 2010) based on the criteria of the American Academy of Sleep Medicine (AASM, American Academy of Sleep Medicine & Iber, 2007). Following the automatic scoring by the algorithm, visual inspection of the results was performed by an expert scorer. KCs were detected automatically using a wavelet-detection algorithm developed by the SIESTA Group. The development and validation procedures of the algorithm are described elsewhere (Parapatics et al. 2015; Schwarz et al. 2017). The detection was carried out in a two-step process; first, the algorithm detects possible KCs via an approach that combines a matched-filtering detection method and a slow-wave detection method (Woertz et al., 2004). First, events that had (a) minimum negative-to-positive peak-to-peak amplitude of 50 mV and (b) a duration between 480 and 1500 ms were detected. Second, all detected events were matched to a prototypical template via wavelet analysis, and via linear discriminant analysis (LDA) to select only real KCs. For our analysis, we considered real KCs to be events that have an LDA score of 0.8 or higher. KCs were detected at C3 and C4, and the events detected at C3 were used for the analysis. Evoked KCs were defined as N2 events that started within the 2 s post-stimulus-onset window. Further details on the sleep architecture related processing are described in Ameen et al., 2022.

### 2.4. PSD estimation

#### 2.4.1. Sleep stage PSDs

For calculating PSDs per sleep stage in iEEG and EEG data, we used Welch’s method (Welch, 1967) with 15 s segments and 50% overlap. We created average PSDs per stages by taking the mean over all segments per epoch then averaging over all epochs.

#### 2.4.2. Frequency range analysis

To compare the performance of the model at different condition of PSD calculation, i.e. using different time windows and frequency ranges for Welch’s PSD calculations, we used four different time windows (5 s, 10 s, 15 s, 20 s) and either four broad frequencies bands (1-30 Hz, 1-45 Hz, 1-60 Hz, 1-75 Hz), and four narrow frequency bands that we selected from our literature search (30-45 Hz, 25-45 Hz, 1-20 Hz, 1-8 Hz). We report the results averaged over all sleep stages of iEEG (Wake, N2, N3, and REM) or EEG (Wake, N1, N2, N3, and REM) data. In our epoch-by-epoch analysis of the exponent, we adopted a methodology similar to what was described in section 2.4.1. This involved calculating the average exponent for each individual epoch within each sleep stage. However, unlike the previous analysis, we did not aggregate the exponent values across all epochs. For the analysis of the exponent over sleep quartiles, we first determined the total count of epochs for each sleep stage. This total was then divided by four to establish the quartiles. Subsequently, we computed the average exponent value for all the epochs contained within each quartile.

#### 2.4.3. Transitions between sleep stages in EEG

To explore the changes in spectral parameters associated with transitions between sleep stages, the night’s EEG data were segmented into 20 s intervals with a moving window of 2 s, resulting in a 90% overlap between adjacent segments. We then applied Welch’s method to calculate the PSD for each of these segments, using the same parameters outlined in section 2.4.1., generating a PSD for each segment. For the analysis of sleep stage transitions, we focused on segments falling within a 120 s timeframe, extending from 60 s before to 60 s after each transition, to capture the spectral dynamics occurring around these changes in sleep stages.

#### 2.4.4. EEG time-resolved analysis

To analyze the temporal aspects of aperiodic activity during auditory stimulation and during KC events, we segmented the EEG data into 10 s epochs centered around the event of interest (stimulus onset or KC onset). The epoched data were inspected and bad epochs containing large-amplitude fluctuations (above 1000 μV) were removed. Subsequently, frequency analysis was performed using a multitaper approach, employing Fast Fourier transformation (FFT)-based convolution between 1 and 45 Hz with 0.5 Hz frequency steps with the number of cycles set to equal 1 s for each frequency. The first and last second of the epoch were discarded to avoid any contamination by edge artifacts. The time-resolved aperiodic estimates were baseline-corrected by subtracting the mean of the values of the 500 ms pre-stimulus-onset window from each data point. For all time-resolved analyses, we used the original sampling rate of the EEG data (250 Hz) except for the KC vs no KC analysis where we downsampled the data to 128 Hz, since KCs were scored on downsampled data. For the comparison between stimulation and no-stimulation, one subject was removed from NREM and another from REM analysis due to poor model fit (as estimated by low R^2^ values). Thus, these analyses were conducted on 16 subjects.

### 2.5. Spectral Parametrization

The specparam toolbox (formerly: ‘FOOOF’, v1.0.0) was used to parametrize iEEG and EEG power spectra (Donoghue et al., 2020). Unless otherwise specified, power spectra were parameterized across the frequency range 1 to 45 Hz. This frequency range was used as a standard, except in cases where we specifically focused on investigating the impact of different frequency ranges. Settings for the algorithm were set as: peak width limits: 1-12 Hz, max number of peaks: 8; minimum peak height: 0; peak threshold: 2. Aperiodic activity was defined according to the formula: 10^b^ x 1 / (k + f^1/x^), where x is the exponent at given frequency range (f), b is the y-axis intercept and k is the knee parameter. The aperiodic mode was set to either ‘fixed’ for a fixed / single-exponent model, i.e. a model that fits a single exponent value to the whole spectrum was used or ‘knee’ when adding a knee parameter to the model, which models a bend in the PSD at a specific frequency after which the exponent changes. The knee frequency is calculated from the knee constant using the formula: knee_freq_ = k^1/x^ (Gao et al., 2020). We calculated the knee frequency from the knee parameter, removing any values that were higher or lower than the mean ± 2 standard deviations. We checked the goodness of fit (R^2^) for all the spectral models. All EEG analyses were performed on electrode Cz, except for the topographical analysis which was done on all 183 EEG electrodes.

### 2.6. Event-related Potential (ERP) analysis

To measure the auditory ERPs of all stimuli as well as the difference between the ERPs of the different stimulus categories (FV vs. UFV and SON vs. UNs), the data were segmented into 10 s epochs centered around stimulus onset. The grand average of all epochs per participant was calculated and baseline correction was performed using the formula (data – mean baseline values) / mean baseline values.

### 2.7. Classification analysis

In order to evaluate the discriminability of different sleep stages based on the estimated aperiodic parameters, classification analysis was done using the scikit-learn toolbox using a linear discriminant analysis (LDA) classifier with K-fold cross-validation (5 splits, 2 repetitions). The number of 30s epochs per stage was equalized and the total number of trials over all stages used for training and testing was equalized between conditions. On average, the number of epochs used per subject was 738.35 ± 567.79 for the iEEG data and 913.82 ± 2.12 for the EEG data. The classifier was trained on 80% of the trials and tested on the remaining 20%. This process was repeated twice and an average of the decoding accuracies was calculated. Chance levels were defined as the reciprocal of the number of alternative outcomes. For instance, in the case of a five-class classification, a chance level of 1/5 or 0.2, denotes the accuracy level that could be achieved only by guessing.

### 2.8. Code

Code for this project was primarily written in the Python programming language (v3.10.6), except for pre-processing of EEG data which was done in Matlab. Preprocessing and segmentation of EEG data were done using EEGLab v14.1.1b in Matlab v. 2019a (Delorme & Makeig, 2004). PSD estimations of iEEG and EEG data were performed in MNE-Python (Gramfort et al., 2013). Spectral parameterization analyses were done using the ‘specparam’ toolbox (https://github.com/fooof-tools/fooof). We deposited the code of this project in the project repository and made it openly available and licensed for reuse at https://github.com/mohamedsameen/Aperiodic_sleep.

### 2.9. Statistical analysis

For comparisons of exponent values and knee frequencies across sleep stages, we performed Friedman chi-square tests and reported Chi-square (X^2^) statistics, p-values, as well as Kendall’s W as a measure of effect size. W was computed by the equation: W = X^2^/N(K-1), where N is the sample size and K is the number of measurements per subject. Kendall’s W uses Cohen’s interpretation guidelines of < 0.3 (small effect), 0.3 - 0.5 (moderate effect) and >= 0.5 (large effect). We performed post-hoc tests, when applicable, via Dunn’s test with Bonferroni’s correction for multiple comparisons and reported the z-values, p-values, and Cliff’s delta (cd) as a measure of effect size which is interpreted using the following values: > 0.33 (small), from 0.33 to 0.474 (medium), and ≥ 0.474 (large). For comparing R^2^ values between broadband and narrowband frequency ranges, we performed a Wilcoxon sign-rank test and reported W as test statistic, p-values and rank biserial correlations (r) as a measure of effect size. Correlations between knee frequency and exponent values were performed either using Pearson’s or Spearman’s according to the results of the normality test and correlation coefficients and p-values were reported.

We measured model performance using the Bayesian Information Criterion (BIC) for Gaussian models (Foygel & Drton, 2010) for the different models, calculated through the equation: BIC = N x log(mse) + np * log(N), where N is the sample size, mse is the mean squared error and np is the number of parameters in the model. Since we fit a maximum of 8 peaks, the number of parameters was equal to: n_peaks * 3 + n_ap_params, where n_peaks depend on the model fit (up to a maximum of 8), and n_ap_params was 2 for the fixed model (offset, exponent), and 3 for the knee model (offset, knee, exponent). A difference in BIC between the two models between 0–2 constitutes ‘weak’ evidence in favor of the model with the smaller BIC, a difference between 2 and 6 constitutes ‘positive’ evidence; and a difference above 6 constitutes ‘strong’ evidence (Raftery, 1995).

Comparisons of classification accuracies against chance levels were done using a permutation t-test where a distribution was generated by shuffling the labels of the sleep stages with those of chance-level then splitting the resulting array into two equal arrays then performing a t-test. This process was repeated 10000 times and the averages of t-values, Bonferroni-corrected p-values, and Cohen’s d effect sizes were calculated. To track the change in aperiodic parameters (exponent and knee frequency) across epochs of sleep stages, a regression analysis was done for every sleep stage using the *statsmodels* toolbox in Python (Seabold & Perktold, 2010) and the regression coefficients (R^2^), the F-statistics, and the p-values were reported. Comparison of time-series data was done using a cluster-based permutation analysis implemented in the *Fieldtrip* toolbox (Maris & Oostenveld, 2007; Oostenveld et al., 2010) in Matlab (v. 2019a) using a two-sided t-test and 5000 permutations. Alpha level was set at 0.025 and we report the sum of t-statistics (∑t) of the cluster, p-values as well as Cohen’s d measured over all possible permutations.

## 3. Results

### 3.1. Model selection for estimating aperiodic activity

In this study, we aimed to broaden the scope of analyzing aperiodic activity during sleep. Our initial step involved a systematic investigation into how the selection of frequency ranges and model forms influences the results. To evaluate the impact of the chosen frequency range on the fitting of the aperiodic model, we manipulated two parameters of the PSD estimation procedure: a) the frequency range the model is fit to, and b) the time window used in Welch’s power estimation, which influences the frequency resolution. For this analysis we fit a fixed exponent model to either broad (Figure 3A) or narrow frequency ranges (Figure 3B) in the iEEG data. The results demonstrated that although all models had consistently high goodness of fit (R^2^), irrespective of frequency range or time window (all > 0.86), R^2^ values were significantly higher for broadband (0.99 ± 0.004) ranges as compared to narrow band (0.95 ± 0.03) ranges (Wilcoxson sign rank test: W = 0, p < 0.001, r = 0.92). Broadband models also showed less variation in R^2^ values between the different frequency ranges and time windows (Figure 3C, Wilcoxson sign rank test: W = 0, p < 0.001, r = 0.96), suggesting that models fit on broadband ranges provide more stable model estimates.

**Figure 3.**
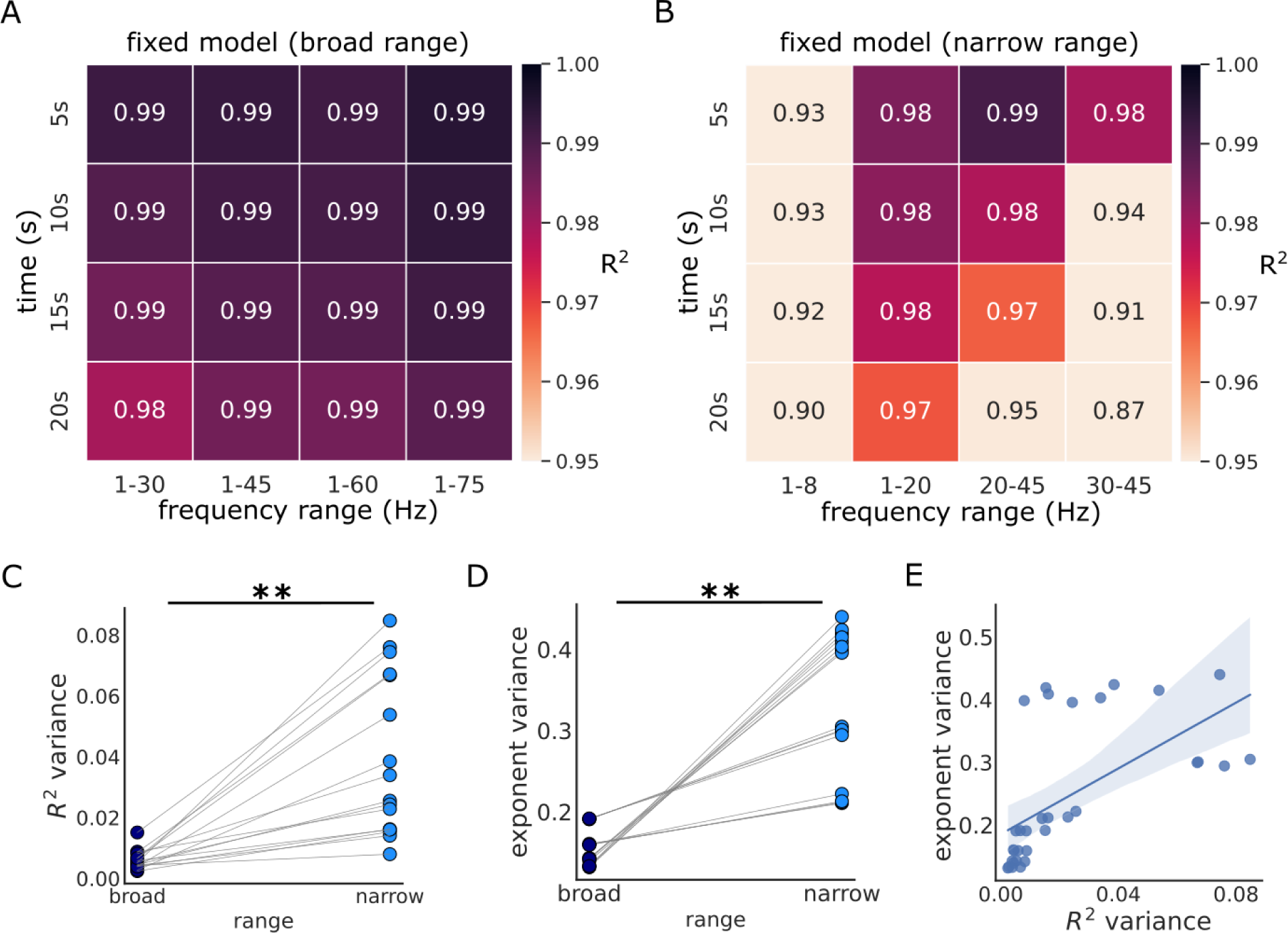
Effects of Frequency Range and Time Window on Model Performance in iEEG Data. A) The R^2^ values, indicating goodness of fit, for models applied to different broad frequency ranges (x-axis) and with different time windows (y-axis). B) The R^2^ values for models using narrow frequency ranges. Note the variability of R^2^ values in comparison to (A). C-D) Models incorporating broadband frequencies demonstrated a marked reduction in variance for C) R^2^ values and D) estimated exponents. E) The relationship between R^2^ variability and exponent variability. A significant positive correlation (r=0.82) was observed between the variances in R^2^ and exponent values. Each point on the graphs symbolizes the value for a particular combination of time window and frequency range, averaged across data from all sleep stages. ** p < 0.001.

Furthermore, the exponent values calculated across various broad and narrow frequency ranges (Suppl. Figure 3-1A-B) exhibited substantially reduced variability with broad ranges relative to those derived from narrow frequency bands (Figure 3D, Wilcoxson sign rank test: W = 0, p < 0.001, r = 1). Additionally, we observed a strong positive correlation between the variance in R^2^ and the variance in the exponent (Figure 3E, Spearman correlation: rho = 0.82, p < 0.001), highlighting the influence of model fits on the variability in exponent values. We observed similar results using the knee model (Suppl. Figure 3-1C-D) as well as in the EEG data (Suppl. Figure 3-2). Collectively, while R^2^ values do not in isolation adjudicate between good and bad models (see for example Figure 2), these results suggest that fitting spectral models with broad frequency range to the data may offer increased reliability.

### 3.2. Stage-specific aperiodic activity in iEEG data

To examine the results of different models for estimating aperiodic parameters in empirical data, we analyzed publicly accessible iEEG sleep data containing epochs from Wake, N2, N3, and REM sleep (von Ellenrieder et al., 2020). In a first step, we visually inspected the PSDs of the different sleep stages, averaged over the recorded cortical regions (Figure 4A). Based on the visible appearance of a knee in the PSDs, we fit a knee model, as seen in example annotated model-fit results (Figure 4B), which better captured the data as compared to using the fixed models (Suppl. Fig. 4-1). Moreover, the single-region PSDs per sleep stage (Figure 4C) demonstrated that when the knees were detected in the average PSD, they were consistently present across the recorded brain areas but varied in their frequencies across stages.

**Figure 4.**
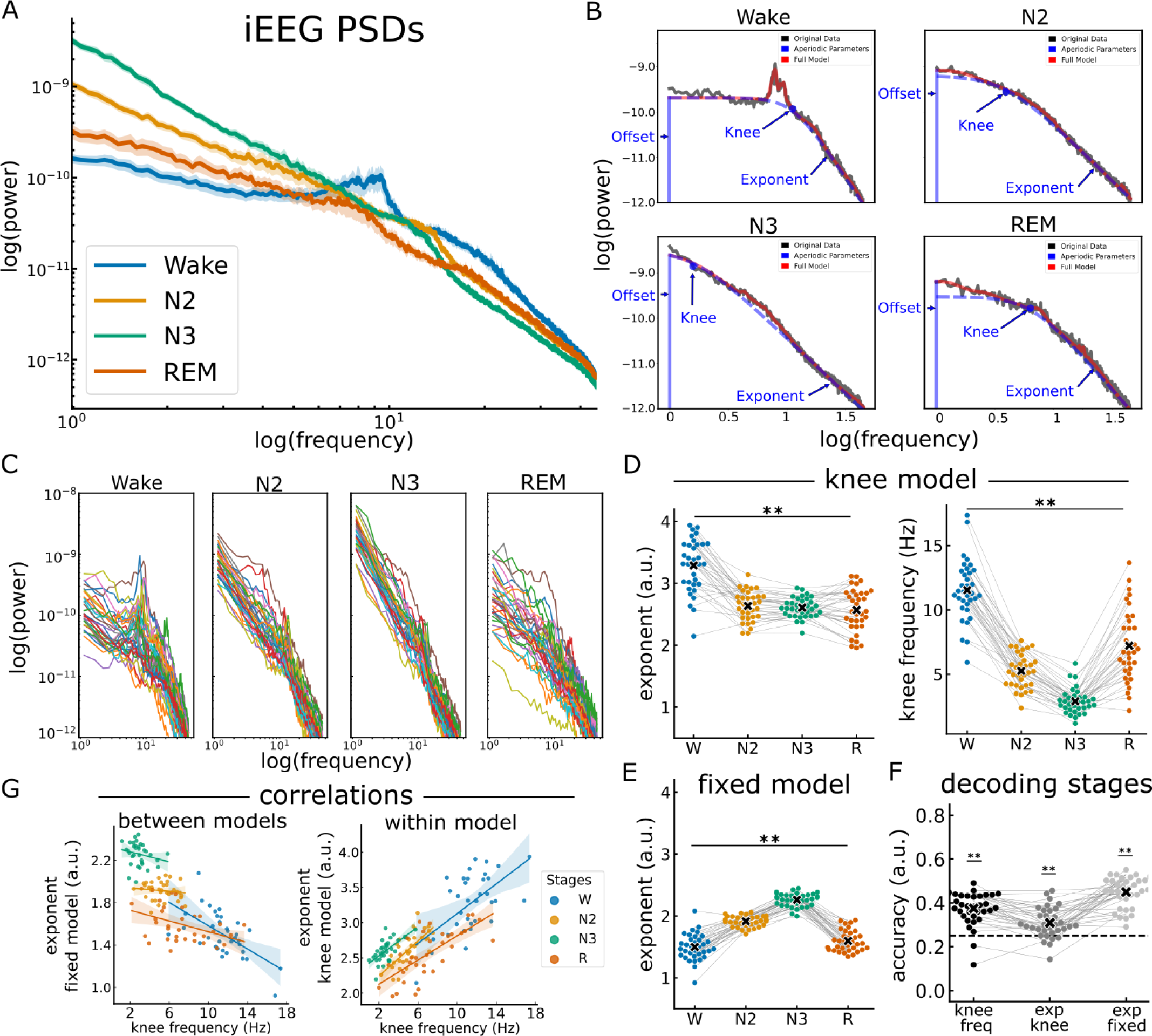
Aperiodic activity in iEEG sleep data. A) Average PSDs of the different sleep stages of the iEEG data. B) An example of the aperiodic fit using the knee broad range model fit in the different sleep stages of the iEEG data, fit on the average PSD of each sleep stage. C) The PSDs of the different sleep stages across 34 cortical regions. D) (Left) The exponent values of the knee model across sleep stages (fit range: 1-45 Hz). Note that the exponent values are different between wakefulness and sleep but not between sleep stages. (Right) The knee frequency across sleep stages. The knee frequency differed significantly between sleep stages. E) The exponent of the fixed model (fit range: 1-45 Hz) was significantly different between stages. F) The classification accuracy between sleep stages using an LDA classifier that uses either the knee frequency, the exponent of the fixed model, or the exponent of the knee model. Note that both the knee frequency and the exponent of the fixed model had significantly above chance classification accuracies. G) Correlations between the knee frequency and the exponents of both the knee and the fixed models. The change in the exponent of the fixed model correlated significantly negative with knee frequency of the knee model. Each dot represents one brain region in the iEEG data. Horizontal lines represent chance level (0.25). ** p < 0.001.

Previous findings have reported that the spectral exponent differs between sleep stages (Höhn et al., 2022; Kozhemiako et al., 2022; Lendner et al., 2020). We sought to replicate these findings using both the knee model and the fixed model. Applying the knee model, the difference in the exponent values between the stages was significant (Figure 4D left; X^2^ = 45.56, p < 0.001, W = 0.45). Post-hoc tests revealed that the exponent differed significantly from wakefulness to: N2 (p_bonf_ < 0.001), N3 (p_bonf_ < 0.001) and REM (p_bonf_ < 0.001). There was no difference, however, between the exponents of the different sleep stages (N2-N3: p_bonf_ = 1, N2-REM: p_bonf_ = 0.98, N3-REM: p_bonf_ = 1), see Table 2 for the detailed statistical results. Notably, the knee frequency exhibited significant stage-dependent differences (Figure 4D right; X^2^ = 89.08, p < 0.001, W = 0.87). The results of the post-hoc tests are reported in Table 3. We also repeated the same analysis using the fixed model and observed significant stage-specific differences in the exponent values (Figure 4E, X^2^ = 90.56, p < 0.001, W = 0.89). The results of the post-hoc tests are reported in Table 4. The model goodness-of-fit values the models are depicted in Suppl. Figure 4-2.

**Table 2.** The statistical results of the post-hoc Dunn’s test to the Friedman test of the difference in exponent values of the knee model between sleep stages in iEEG. z: z-value p: Bonferroni-corrected p-values, cd: Cliff’s delta effect size.

|  | Pairwise | z | p | cd |
| --- | --- | --- | --- | --- |
| 0 | Wake vs N2 | 5.23 | > 0.001 | 0.813 |
| 1 | Wake vs N3 | 5.694 | > 0.001 | 0.86 |
| 2 | Wake vs REM | 5.816 | > 0.001 | 0.808 |
| 3 | N2 vs N3 | > 0.001 | 1.0 | 0.073 |
| 4 | N2 vs REM | > 0.001 | 1.0 | 0.1 |
| 5 | N3 vs REM | > 0.001 | 1.0 | 0.045 |

**Table 3.** The statistical results of the post-hoc Dunn’s test to the Friedman test of the difference in knee frequencies between sleep stages in iEEG. z: z-value p: Bonferroni-corrected p-values, cd: Cliff’s delta effect size.

|  | Pairwise | z | p | cd |
| --- | --- | --- | --- | --- |
| 0 | Wake vs N2 | 5.23 | > 0.001 | 0.981 |
| 1 | Wake vs N3 | 5.694 | > 0.001 | 1.0 |
| 2 | Wake vs REM | 5.816 | 0.001 | 0.753 |
| 3 | N2 vs N3 | > 0.001 | 0.001 | 0.856 |
| 4 | N2 vs REM | > 0.001 | 0.23 | -0.448 |
| 5 | N3 vs REM | > 0.001 | > 0.001 | -0.896 |

**Table 4.** The statistical results of the post-hoc Dunn’s test to the Friedman test of the difference in exponent values of the fixed model between sleep stages in iEEG. z: z-value p: Bonferroni-corrected p-values, cd: Cliff’s delta effect size.

|  | Pairwise | z | p | cd |
| --- | --- | --- | --- | --- |
| 0 | Wake vs N2 | 5.041 | > 0.001 | -0.922 |
| 1 | Wake vs N3 | 8.21 | > 0.001 | -0.997 |
| 2 | Wake vs REM | > 0.001 | 1.0 | -0.292 |
| 3 | N2 vs N3 | 3.403 | 0.001 | -0.986 |
| 4 | N2 vs REM | 3.833 | > 0.001 | 0.874 |
| 5 | N3 vs REM | 7.896 | > 0.001 | 1.0 |

To test which parameter is a better predictor of the difference between sleep stages, we employed an LDA classifier trained on the knee frequency, the exponent of the knee model, or the exponent of the fixed model (Figure 4F). The classification results revealed significantly above chance-level classification accuracy using the knee frequency (t(33) = 6.77, p_bonf_ < 0.001, d = 2.33), the exponent of the knee model (t(33) = 3.55, p_bonf_ < 0.001, d = 1.22) and the exponent of the fixed model (t(33) = 11.93, p_bonf_ < 0.001, d = 4.11). Comparing the accuracies of these three parameters demonstrated a significant difference between the groups (X^2^ = 35.38, p < 0.001, W = 0.52). The accuracy using the knee frequency was significantly higher than that of the exponent of the knee model (z(33) = 4.57, p_bonf_ < 0.001, cd = 0.8), but lower than that of the exponent of the fixed model (z(33) = 3.07, p_bonf_ = 0.006, cd = -0.59). The accuracy using the exponent of the fixed model was higher than that of the exponent of the knee model (z(33) = 7.64, p_bonf_ < 0.01, cd = 0.92).

Given this pattern of multiple aperiodic parameters relating to sleep stages, we next sought to explore the potential relationship between these parameters. To do so, we computed correlations between the knee frequency and the exponent estimates. Between models – comparing the knee frequency of the knee model and the exponent of the fixed model – there was a significant negative correlation (Figure 4G left; rho = -0.82, p < 0.001). However, this relationship is reversed when examining the exponent values resulting from the knee model. Specifically, the correlation between the knee frequency and the exponent of the knee model was positive and significant (within-model correlation: Figure 4G right; rho = 0.69, p < 0.001). These findings highlight a potential interdependence between the knee frequency and the exponent – suggesting that if there is a knee in the signal, when measuring the exponent of the fixed model these results may relate to changes in the knee frequency. In the context of sleep data, this suggests that previously reported differences in the exponent might actually reflect differences in the knee frequency between sleep stages, as seen here when explicitly measuring knee models.

### 3.3. Stage-dependent patterns in EEG aperiodic activity

Based on these results in the iEEG analysis, we next examined sleep-related aperiodic activity in a full night recording of EEG, as most previous results are in extracranial data. Starting again with a visual inspection, we saw that the PSDs across distinct sleep stages revealed differences in the exponent across stages, as well as a stage-dependent knee (Figure 5A). Specifically, we observed a prominent knee during REM sleep that appears to be attenuated in wakefulness, N1 and N2, and looks to be absent during N3. Figure 5B illustrates PSDs for different sleep stages, across participants, emphasizing the consistency of stage differences across all regions, and highlighting the consistency of the knee parameter in REM sleep.

**Figure 5.**
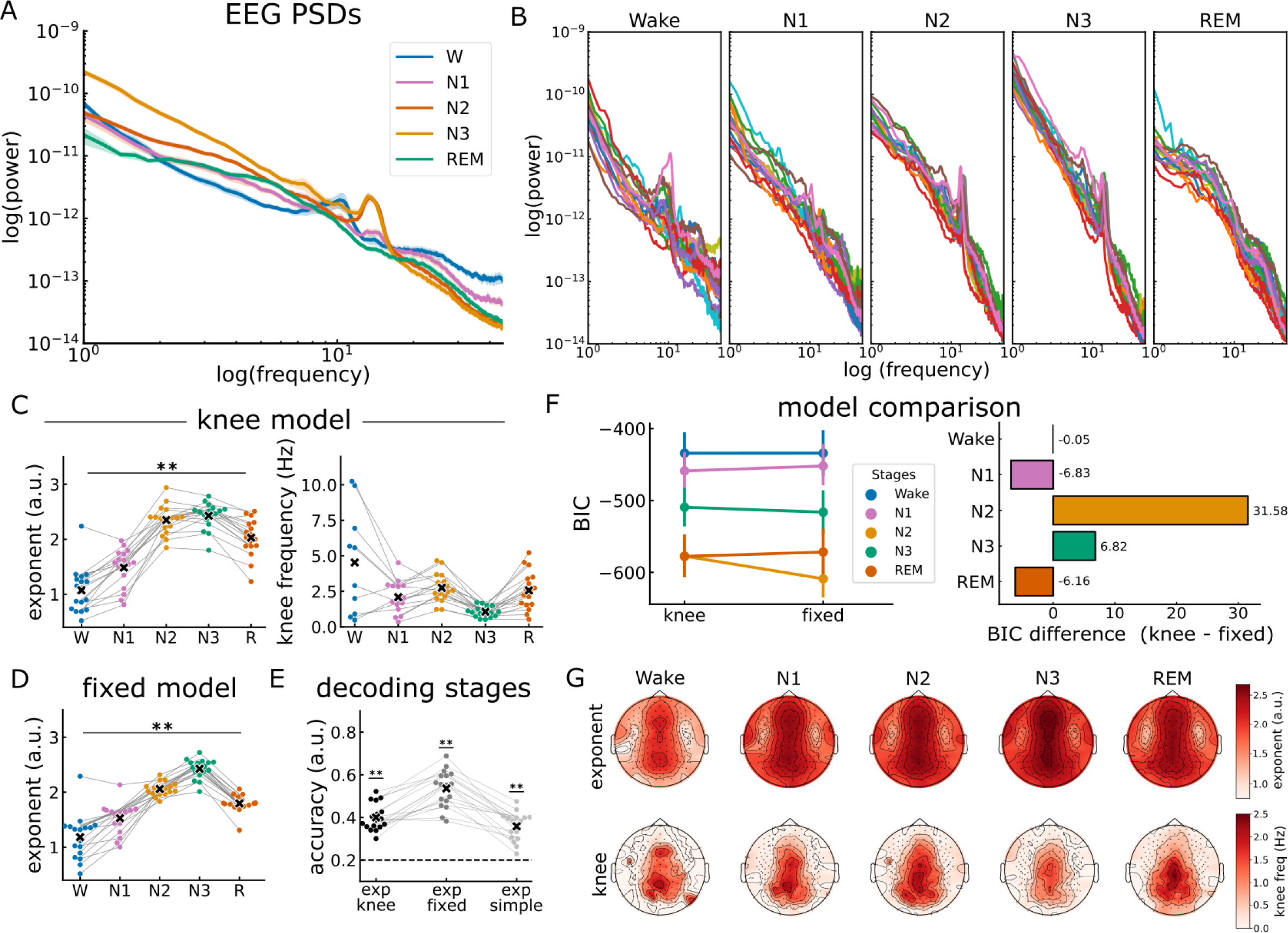
Aperiodic activity in EEG sleep data. A) PSDs of the different sleep stages calculated over electrode Cz. B) The PSDs of the different stages per subject measured at electrode Cz. Note the stage-dependent observation of the ‘knee’ in the PSD. C) (left) Exponent values and (right) the knee frequency values of the knee model (fit range: 1-45 Hz)) across sleep stages. D) Exponent values of the fixed model across sleep stages (fit range: 1-45 Hz). The exponents of both the knee model and the fixed model significantly differed between sleep stages, while the knee frequency did not. E) The classification accuracy between the EEG exponent of the different sleep stages using an LDA classifier trained on the fixed model, the knee model, and the fixed model with the narrow band (30-45 Hz). Regardless of the frequency range used, the classifier was able to use the exponent to classify between the stages with an above chance (0.2) accuracy. F) Comparison of model performance between the fixed model and the knee model across sleep stages using the BIC metric. BIC was significantly lower (better fit) for the knee model as compared to the fixed model in REM sleep (-6.15), and N1 (-6.83), but was significantly higher (worse) for N2 (31.58) and N3 (6.82). For wakefulness, however, the models did not significantly differ (-0.05). G) Topographical maps depicting the spatial distribution of the exponent of the knee model (top) and the knee frequency (bottom) across the scalp. Note the consistency of the central topography of the exponent and the knee frequency across stages. Each dot represents one participant of the EEG dataset. Horizontal lines represent chance level (0.2). * p < 0.05, ** p < 0.001.

We next fit spectral models to the data, with the goal of replicating the analyses from the iEEG data, in order to evaluate the effectiveness of the exponent and knee parameters in differentiating between sleep stages in EEG data (Figure 5C-D). When applying the knee model (fit range: 1-45 Hz), significant differences were observed in the exponent across stages (Figure 5C left; X^2^ = 58.21, p < 0.001, W = 0.86). The detailed outcomes of the post-hoc tests can be found in Table 5. However, in contrast to the iEEG data, the knee frequency did not exhibit significant differences between stages (Figure 5C right, X^2^ = 30.01, p < 0.001, W = 0.44), see Table 6 for post-hoc statistical results. When applying a fixed model (fit range 1-45 Hz), the exponent also showed a significant difference between sleep stages (Fig. 5D; X^2^ = 61.93, p < 0.001, W = 0.91). The results of the post-hocs test are detailed in Table 7. The R^2^ of the various models for different sleep stages is depicted in Suppl. Figure 5-1A-B.

**Table 5.** The statistical results of the post-hoc Dunn’s test to the Friedman test of the difference in exponent values of the knee model between sleep stages in EEG. z: z-value p: Bonferroni-corrected p-values, cd: Cliff’s delta effect size.

|  | Pairwise | z | p | cd |
| --- | --- | --- | --- | --- |
| 0 | Wake vs N1 | > 0.001 | 1.0 | -0.619 |
| 1 | Wake vs N2 | 5.232 | > 0.001 | -0.958 |
| 2 | Wake vs N3 | 6.068 | > 0.001 | -0.979 |
| 3 | Wake vs REM | 3.299 | 0.001 | -0.889 |
| 4 | N1 vs N2 | 3.768 | > 0.001 | -0.986 |
| 5 | N1 vs N3 | 4.638 | > 0.001 | -0.986 |
| 6 | N1 vs REM | 1.628 | 0.103 | -0.765 |
| 7 | N2 vs N3 | > 0.001 | 1.0 | -0.287 |
| 8 | N2 vs REM | 0.239 | 0.811 | 0.516 |
| 9 | N3 vs REM | 1.579 | 0.114 | 0.702 |

**Table 6.** The statistical results of the post-hoc Dunn’s test to the Friedman test of the difference in knee frequency between sleep stages in EEG. z: z-value p: Bonferroni-corrected p-values, cd: Cliff’s delta effect size.

|  | Pairwise | z | p | cd |
| --- | --- | --- | --- | --- |
| 0 | Wake vs N1 | > 0.001 | 1.0 | 0.294 |
| 1 | Wake vs N2 | > 0.001 | 1.0 | 1.0 |
| 2 | Wake vs N3 | 2.398 | 0.017 | 1.0 |
| 3 | Wake vs REM | > 0.001 | 1.0 | 1.0 |
| 4 | N1 vs N2 | > 0.001 | 1.0 | -0.1 |
| 5 | N1 vs N3 | 1.142 | 0.253 | 0.626 |
| 6 | N1 vs REM | > 0.001 | 1.0 | 0.024 |
| 7 | N2 vs N3 | 3.097 | 0.002 | 0.924 |
| 8 | N2 vs REM | > 0.001 | 1.0 | 0.128 |
| 9 | N3 vs REM | 2.337 | 0.019 | -0.73 |

**Table 7.** The statistical results of the post-hoc Dunn’s test to the Friedman test of the difference in exponent values of the fixed model between sleep stages in EEG. z: z-value p: Bonferroni-corrected p-values, cd: Cliff’s delta effect size.

|  | Pairwise | z | p | cd |
| --- | --- | --- | --- | --- |
| 0 | Wake vs N1 | > 0.001 | 1.0 | -0.619 |
| 1 | Wake vs N2 | 4.683 | > 0.001 | -0.889 |
| 2 | Wake vs N3 | 6.863 | > 0.001 | -0.979 |
| 3 | Wake vs REM | 2.294 | 0.022 | -0.827 |
| 4 | N1 vs N2 | 3.275 | 0.001 | -0.91 |
| 5 | N1 vs N3 | 5.537 | > 0.001 | -0.993 |
| 6 | N1 vs REM | 0.374 | 0.708 | -0.779 |
| 7 | N2 vs N3 | 0.835 | 0.404 | -0.875 |
| 8 | N2 vs REM | 0.871 | 0.384 | 0.875 |
| 9 | N3 vs REM | 3.555 | > 0.001 | 0.993 |

To investigate how aperiodic parameters vary across sleep stages, we examined the classification accuracies using the different exponent estimates, including the exponent of the a) knee model, b) fixed model broad range, and c) fixed model narrow range (30-45 Hz), which was added to compare to previous reports. The results showed that the exponents of all models performed significantly higher than chance levels (exp_knee: t(16) = 12.98, p_bonf_ < 0.001, d = 4.45, exp_fix_broad: t(16) = 10.81, p_bonf_ < 0.001, d = 3.71, exp_fix_narrow: t(16) = 9.81, p_bonf_ < 0.001, d = 3.37). Comparing the accuracies of these three parameters demonstrated a significant difference between the accuracy values from the different models (Figure 5E; X^2^ = 24.39, p < 0.001, W = 0.72). Post-hoc tests revealed that the exponent of the fixed model (broad range) performed significantly higher than that of the knee model (z(16) = 3, p_bonf_ =b 0.009, d = 0.66) and the exponent fit over the 30-45 (narrow) range (z(16) = 4.11, p_bonf_ < 0.001, d = 0.75). There was no difference between the exponent of the knee model and the narrow range (z(16) = 1.15, p_bonf_ = 0.74, d = 0.3).

To assess and compare the performance of the different models, we used the bayesian information criterion (BIC) index for gaussian models (Fig. 5F). The BIC is a criterion for model selection that is based on the likelihood function and takes into account the number of parameters in the model. This approach allows for a more comprehensive evaluation, as it not only considers the fit of the model to the data but also penalizes models with a larger number of parameters. The results revealed a sleep-stage-dependent performance for these models. Specifically, BIC was lower for the knee model as compared to the fixed model in REM sleep (-6.15), N1 (-6.83), suggesting a better fit using the knee model in these stages. Conversely, BIC was higher for the knee model as compared to the fixed model for N2 (31.58) and N3 (6.82) suggesting a better fit using the fixed model. For wakefulness however, BIC was not significantly different between the models (BIC = -0.05). Overall, these analyses of model forms and aperiodic parameters relationships to sleep stages suggest a more nuanced pattern than in the intracranial data – while there are observable knees in the data, with differences across stages, the variability of this parameter is such that the knee frequency is less predictive of different sleep stages.

We also examined the aperiodic parameters across the scalp, whereby the topographies for the exponent of the knee model and the knee frequency across sleep stages show a consistent spatial pattern of the largest exponent and knee frequencies around the central and posterior electrodes, with this topography being consistent across sleep stages (Figure 5G; see Suppl. Figure 5-1C for the topography of the fixed model).

### 3.4. Time-resolved EEG exponent tracks changes in sleep architecture

Having established differences in aperiodic parameters across different sleep stages, our next aim was to examine the temporal dynamics of aperiodic activity during sleep (Figure 6A). We started by investigating the influence of time on the exponent during different sleep stages. To do so, we measured the exponent for each epoch within every sleep stage then conducted regression analyses for each stage separately (Figure 6B). The findings revealed distinct temporal patterns in the exponent values across different sleep stages. During wakefulness, there was a notable increase in the exponent value as the night advanced. A significant portion of this increase could be attributed to the passage of time (*R*^2^ = 0.83, F(1,11) = 55.1, p < 0.001). Conversely, we observed a significant effect of time on the decrease in the exponents during N3 and REM (N3: *R*^2^ = 0.26, F(1,129) = 46.66, p < 0.001, REM: *R*^2^ = 0.07, F(1,90) = 7.24, p = 0.008). For N1, although there was an upward trend in the exponent values, the change was not statistically significant (*R*^2^ = 0.13, F(1,12) = 1.76, p = 0.21) while during N2 sleep, the exponent values remained relatively stable over time with no significant change (*R*^2^ = 0.002, F(1,190) = 0.3, p = 0.58).

**Figure 6.**
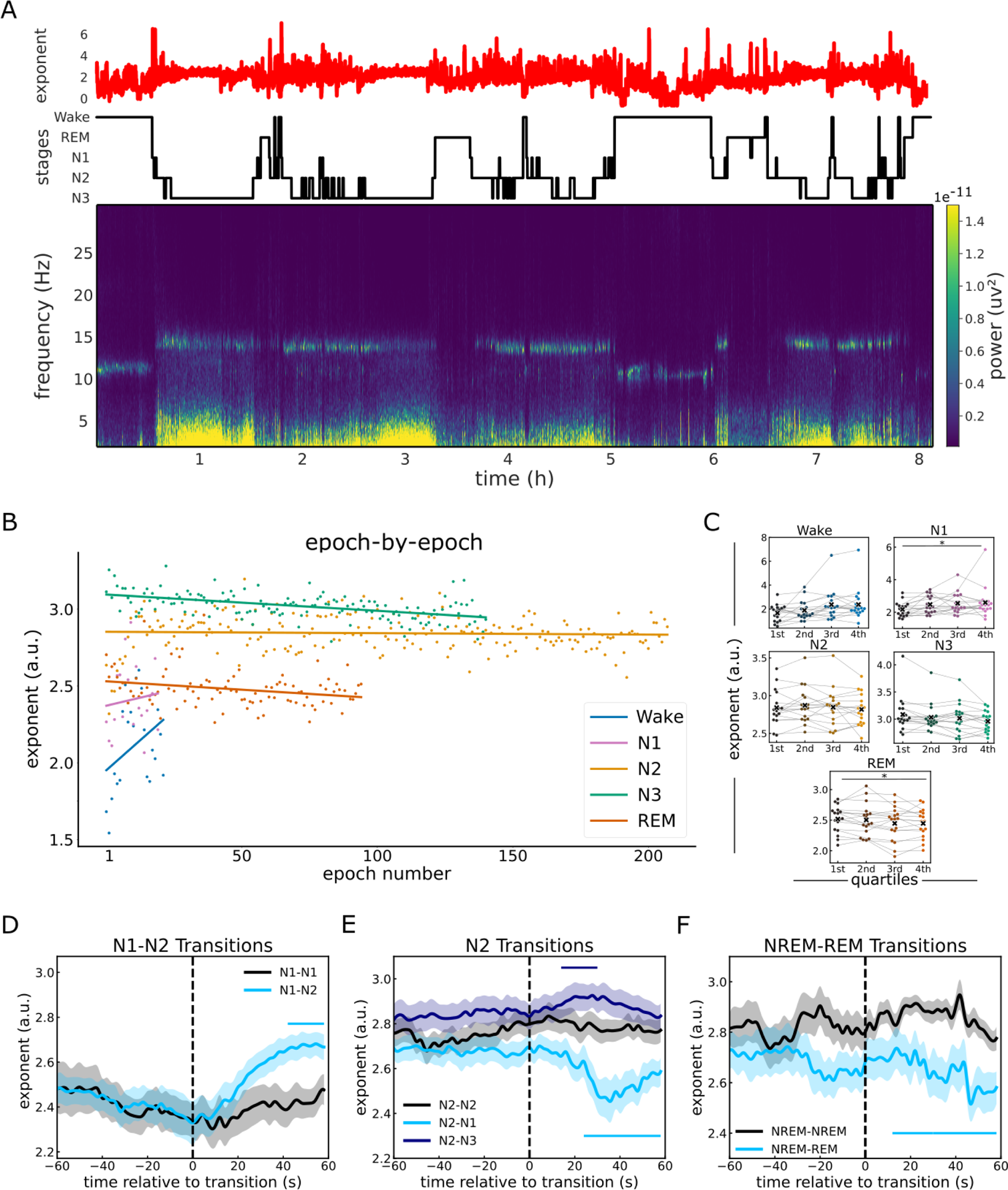
EEG exponent tracks the dynamics of sleep macrostructure. A) Temporally resolved estimates of the spectral exponent, overlaid on top of the sleep staging and spectral plot for the entire night, taken from an example subject. B) Epoch-by-epoch exponent values from across the night for each sleep stage. Notably, Wake and N1 show an increase in the exponent as the night progresses while N3 and REM show a decrease, and N2 shows no significant change. C) The stage-specific exponent values split across the quartiles of the night, where each dot represents a subject. The exponent shows no significant differences across the quartiles, except for N1 where it increases and REM sleep where it decreases significantly across the night. D-F) Time-resolved estimates of the spectral exponent during sleep stage transitions. Each transition is compared to a control period of adjacent epochs when no change in sleep stage occurred. D) Sleep stage transition from N1, showing an increase in the exponent in the transition from N1 to N2 sleep stages, which is significant starting 24 s after transition. E) Sleep stage transitions from N2 to either N1 or N3. Post-transition from N2 to N1, the exponent significantly decreased after 18 s. Conversely, transitioning from N2 to N3 led to an increase in the exponent, this change was significantly different from the period of uninterrupted N2 sleep in a very short time window (14s-30s). F) Transitions from NREM to REM sleep, showing a significant decrease in exponent starting after +18 s from the transition. All exponent values in this analysis reflect the exponent from a knee model (fit range: 1-45 Hz). Vertical dashed lines in D-F indicate the transitions between stages. Horizontal lines indicate significant clusters.

To further examine the fluctuations in aperiodic activity over the course of the night, we assessed the changes in the exponent across quartiles by dividing epochs for each sleep stage into quartiles and comparing exponent measures across these quartiles (Figure 6C). In this analysis, the exponent during REM decreased significantly over quartiles of the night (X^2^ = 13.31, p = 0.004, W = 0.26), while the exponent of N1 increased significantly across the night (X^2^ = 8.65, p = 0.001, W = 0.17). There was no significant difference for Wake (X^2^ = 3.21, p = 0.36, W = 0.06), N2 (X^2^ = 2.29, p = 0.51, W = 0.04), or N3 (X^2^ = 6.04, p = 0.11, W = 0.12).

Next, we sought to evaluate the temporal precision of changes in aperiodic activity. Thus, we analyzed time-resolved exponent values during transitions between sleep stages, as identified by our sleep scoring algorithm. To gain insights into how the exponent values change during these transitions, we compared them to a baseline period of equal length where no change in sleep staging occurred. We specifically focused on the most prevalent transitions during sleep, including N1 to N2, N2 to N1, and N2 to N3, as indicated by the transition matrix (Suppl. Figure 6-1). Comparing the transitions from N1 to N2 against a baseline of continuous N1 (Figure 6D), we found that the exponent increased shortly after the transition from N1 to N2 reaching a significant difference to that of the baseline starting at 24s following the transition (28.71 ± 10.02 trials - 24-60s: ∑t(16) = 87.82, p < 0.001, d = 1.38). Similarly, when examining transitions from N2 to either N3 or N1 (Figure 6E), the results revealed a significant decrease in the exponent 18 s after transition from N2 to N1 (17.59 ± 7.1 trials - ∑t(16) = 164.24, p < 0.001, d = 1.58), and a significant increase in the exponent 14 s after the transition from N2 to N3 (22.18 ± 6.81 trials - 14-30s: ∑t(16) = 33.19, p = 0.003, d = 1.38). Furthermore, we sought to examine a pivotal transition in sleep that marks a significant change in brain activity: the shift from NREM sleep to REM sleep (Figure 6F). For this purpose, we analyzed the exponent during the transition from either N2 or N3 sleep to REM sleep, contrasting it with a baseline period of uninterrupted NREM sleep. The results demonstrate a significant decrease in the exponent during the NREM-to-REM transition as compared to the NREM baseline (4 ± 1.53 trials - 16 - 60s: ∑t(16) = 73.39, p = 0.002, d = 1.12). Overall, these results are consistent with the expected changes in aperiodic exponent from non-time-resolved analyses, and emphasize the temporal precision of these changes in sleep stage. We also repeated the time-resolved analyses of sleep stage transitions using both the exponent of the fixed model and the knee frequency (Suppl. Figures 6-2 and 6-3). The results are broadly consistent, though less reliable and temporally precise as the exponent of the knee model, suggesting that the knee model may be well posed for detecting variations during the transitions between sleep stages.

### 3.5. Selective EEG exponent responses to auditory stimuli during sleep

Next, we aimed at investigating how the exponent responded to external events by analyzing evoked responses to auditory stimuli. To do so, we first computed and analyzed time-resolved measures of aperiodic activity time locked to auditory stimuli, as compared to baseline periods of equal length where no stimuli were presented. During NREM (N2 and N3) sleep, we observed a significant increase in the exponent following stimulus presentation (Figure 7A, -0.08s to 1.12s: ∑t(15) = 1029.54, p < 0.001, d = 1.94). It’s noteworthy that the average duration of these stimuli was 808 ms, suggesting that the transient exponent dynamics match the time-course of the stimulus presentation. In contrast, during REM sleep, there was no observable change in the exponent following the presentation of stimuli (Suppl. Figure 7-1A). To ensure the validity of the observed changes in the exponent values and to confirm that these changes were genuine and not a result of poor model performance, we assessed the R^2^ values around the time of stimulus presentation. The R^2^ values were found to be high (all R^2^ > 0.99) and stable across time (Suppl. Figure 7-1B-C).

**Figure 7.**
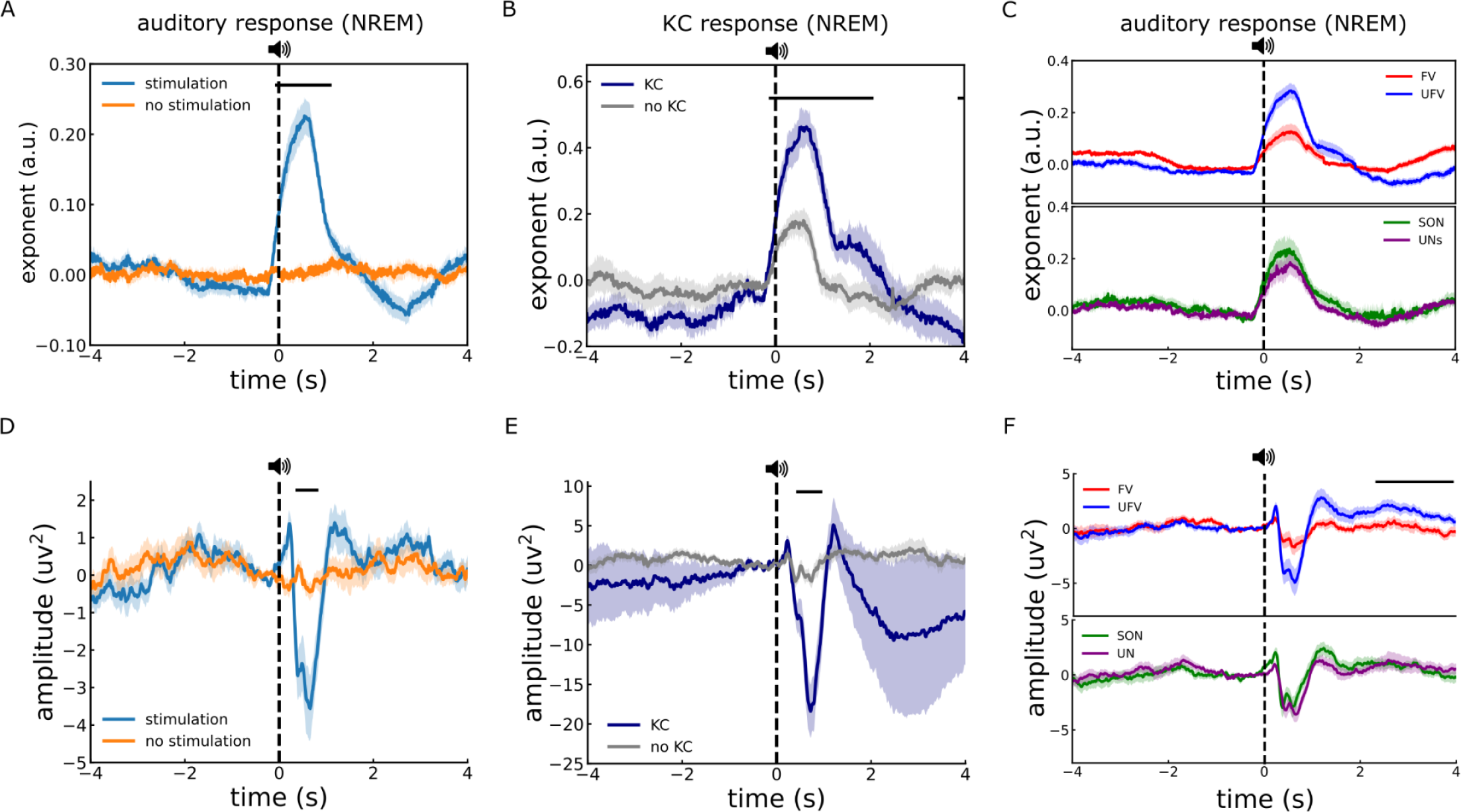
Auditory-evoked exponent and ERP responses during sleep. A) Time-resolved estimates of aperiodic exponent in response to auditory stimuli (blue) compared to data segments with no stimuli (orange) during NREM sleep (N2 and N3), showing a transient, time-locked increase in exponent values during stimulus presentation. B) Comparison of auditory-evoked exponent changes for trials with and without elicited K-complexes during N2. C) Auditory-evoked exponent responses split by auditory stimuli, comparing unfamiliar voices (UFV) and familiar voices (FV), as well as one’s own name (SON) and unfamiliar names (UNs). Aperiodic exponent responses differ by voice familiarity, but not by stimulus content. D) As in A for ERP responses, showing temporal responses to sounds during NREM sleep. E) As in B for ERP responses, comparing stimulus trials with and without evoked K-complexes. F) As in C for ERPs, comparing responses across stimulus conditions, showing ERP responses also differentiate the familiarity of voice. Across all panels, the onset of the stimulus is marked by dashed vertical lines, and clusters representing significant differences between plotted measures are indicated by solid horizontal lines.

We noted that the aperiodic responses we observed are similar to the typical auditory-evoked KC responses that have also been reported during NREM sleep (Ameen et al., 2022). To evaluate how these transient aperiodic responses related to KC activity, we compared the changes in the exponent in trials where a KC was elicited with those where no KCs were observed (Figure 7C). The results indicated that trials in which a KC was evoked showed a significantly larger change in the exponent compared to trials without a KC (-0.14 to 2.07; ∑t(16) = 789.86, p < 0.001, d = 1.99). This suggests a possible association between the aperiodic evoked response and the auditory-evoked KC. Indeed, when we conducted a comparison between the PSDs at the peak of the exponent responses when KCs were elicited and when no KCs were elicited, we observed a broadband power difference suggestive of an exponent shift (Suppl. Fig. 7-2, 1-13 Hz: ∑t(16) = 234.54, p < 0.001, d = 1.23; 15-45 Hz: ∑t(16) = 103.06, p = 0.003, d = 1.35). Thus an aperiodic shift in neural activity might be underlying the KC – though we still observed exponent responses in trials in which no KC was detected, suggesting that these two phenomena are not entirely overlapping.

We next examined whether these evoked responses of the aperiodic exponent are sensitive to different types of stimuli. Our investigation encompassed a range of stimulus categories, particularly contrasting familiar voices (FV) with unfamiliar voices (UFV), and the subject’s own name (SON) against unfamiliar names (UNs) during both NREM and REM sleep (Figure 7D). During NREM sleep, the analysis indicated a significantly larger response of the exponent following the presentation of UFVs as compared to FVs (Figure 7D-Top, -0.14s to 1.03s: ∑t(16) = 577.26, p = 0.003, d = 1.19). Subsequently, the exponent for UFVs decreased to be significantly lower than FVs (2.44 s to 4 s: ∑t(16) = 793.48, p = 0.001, d = 1.06). No discernible difference was found when comparing the exponent’s response to the SON and UNs (Figure 7D-Bottom). During REM sleep, the responses did not significantly differ between either the different voices or the names (Suppl. Figure 7-1E).

Finally, we also examined time-domain responses by examining the auditory ERPs for all auditory stimuli, as well as for the different categories of stimuli. The analysis revealed negative peaks in response to auditory stimuli during NREM sleep (Figure 7E, ∑t(16) = -249.72, p = 0.003, d = 0.9), but no difference to no-stimulation periods in REM sleep (Suppl. Figure 7-1D). The ERPs exhibited a stronger negative peak in trials that triggered a KC (Figure 7G, 0.41 s - 0.96 s: ∑t(16) = -249.72, p = 0.003, d = -1.03). Further, while we observed a negative peak following UFV presentations compared to FVs between 0.45 s and 0.72 s, it lacked statistical significance (Figure 7H-Top; ∑t(16) = -151.97, p = 0.12, d = 0.56). However, a significant positive deflection was noted following UFVs in the duration from 2.23 s to 3.96 s (∑t(16) = 1293.46, p < 0.001, d = 0.89). No significant variations were observed in the responses to different names (Figure 7H-Bottom). Last but not least, ERPs during REM showed no difference between either voices or names (Suppl. Fig. 7-1F). Taken together, our findings once more point to a potential overlap between measures of ERPs, KCs, and the aperiodic exponent in neural activity during sleep, though with some notable distinctions between them. One important distinction is that we observe stimulus-related responses in the exponent when no KCs are detected, when there is also no clear ERP response.

## 4. Discussion

Recent investigations of electrophysiological data have established that aperiodic neural activity offers novel information about brain function, beyond what is available through the use of conventional approaches that primarily focused on periodic, oscillatory aspects (Donoghue & Watrous, 2023; He, 2014). This includes research establishing that aperiodic activity systematically varies across the sleep cycle (Feinberg et al., 1984; Lendner et al. 2020; Höhn et al. 2023; Kozhemiako et al. 2022; Miskovic et al., 2019). In this work, we sought to extend the investigation of aperiodic activity, by examining it across modalities (EEG and iEEG), frequency ranges, model forms, and across time. By doing so, we were able to replicate previous findings of variations of aperiodic activity during sleep, while extending these results to show how multiple aperiodic features show variations across sleep with temporal dynamics that track sleep-stage transitions and neural responses to external stimulation. Overall, these findings suggest that by improving our approaches for measuring aperiodic activity, we can capture aperiodic fluctuations that reflect both macro- and micro-structures of sleep, offering a more comprehensive view of sleep dynamics.

In evaluating the previous literature on aperiodic activity during sleep, we noted that previous studies have typically employed a variety of different frequency ranges to estimate aperiodic activity, with no clear consensus. We therefore investigated the influence of varying the frequency range across different frequency ranges. Overall we found there was high model fit (R^2^ > 0.86) across all frequency ranges, however when using narrow frequency ranges there was a trend for lower model fit quality, more dependency on the length of the time window, and higher variability in model fit quality and parameter results. This suggests that while different frequency ranges can be validly examined, choosing broader ranges may offer more stable estimates. Indeed, the use of broader frequency ranges led to reduced variance in the derived parameters, suggesting increased reliability. Thus, we suggest that future studies of aperiodic activity during sleep should clearly indicate which fit ranges are used, consider fitting broader ranges, where appropriate, and, where possible, include sensitivity analysis examining the dependency of findings on the fit range.

Another notable aspect of the previous literature is the use of ‘fixed’ exponent models, which estimate the spectral exponent by fitting the equivalent of a straight line in the log-log space within a narrow frequency range, typically, between 30 - 50 Hz (Gao et al., 2017; Lender et al., 2020). While this approach has proven effective in distinguishing between different sleep stages, it does not capture the breadth of aperiodic activity, which by definition extends across all frequencies. Notably, fitting a single exponent to a narrow band range ignores the potential presence of a ‘knee’ or bend in the PSD, after which the exponent changes (Gao et al., 2020; Miller et al., 2009). By examining broader frequency ranges, analyses can capture more variance in the data, including of slower frequencies which are particularly prevalent during sleep, and can also explicitly measure the knee, allowing for examination of the knee frequency and what it reflects. Notably, selection of frequency range and model forms should be considered together – for example, examining whether fitting a broader frequency range would require fitting a knee parameter – as qualitative differences in the structure of the data can impact model fitting, even if quantitative measures of goodness-of-fit measures such as R^2^ are similar.

By applying a spectral parameterization approach that examined broad frequency ranges and fit an aperiodic model with a knee, we demonstrate differences in aperiodic activity between sleep stages beyond what can be seen with a single-exponent (fixed) model. Crucially, in the intracranial data, our findings demonstrate that when fitting a knee model, it is the knee frequency that is most effective at distinguishing between sleep stages. This suggests that in this data a prominent change across sleep stages is primarily in the knee frequency, whereby this can look like an exponent change if only analyzing a narrow frequency range using a single-exponent model. Further supporting this, we observed a negative correlation between the knee frequency (in the broad range knee model) and the exponent of the fixed model. Overall, while a single exponent fit within a narrow band frequency may suffice if the primary goal is to differentiate between sleep stages, the underlying changes in the data may be better captured by fitting more complex models over broader frequency ranges.

Notably, the presence of a knee in electrophysiological signals is not universal, particularly in EEG in which very little research has sought to measure a knee as it is usually assumed to be absent. In the EEG dataset examined here, a knee is most prominently visible during REM sleep, while being somewhat visible in Wake and N1, and seemingly absent during NREM stages N2 and N3. However, the seemingly variable presence of the knee in the EEG data appears to be different from what is seen in the intracranial data, and this variability introduces a model-selection challenge. Notably no single model definition is best across all sleep stages in EEG, as the knee model demonstrated a better fit in N1 and REM, while the fixed model outperformed in N2 and N3. This underscores that while both models are suitable for a qualitative differentiation between sleep stages, an informed model-selection decision is crucial for accurately and quantitatively investigating the specific neural activity patterns within each stage.

In terms of interpretations of aperiodic neural activity, previous investigations that largely focused on changes in the aperiodic exponent have typically interpreted changes in this parameter in relation to its putative relation to excitation-inhibition (E-I) balance, whereby a steeper slope signifies an increase in inhibition, and conversely, a shallower slope indicates an increase in excitation (Gao et al., 2017; Chini et al., 2022). The pattern of changes in previous studies is consistent with a general shift in E-I balance across the sleep wake cycle with a steeper slope / more inhibition during NREM (as compared to wake), and a flatter slope / more excitation during REM (as compared to NREM) (Lendner et al., 2020; 2023). This pattern of results is broadly consistent with invasive work in animal models reporting increased inhibition (Birdi et al., 2020) and decreased pyramidal activity (Niethard et al., 2016) during sleep. Our findings here are at least partially consistent with these findings, whereby across all the different model settings this general pattern of exponent differences between Wake, NREM, and REM stages is observed. However, beyond the exponent, measuring aperiodic activity in this study also demonstrated prominent changes in the knee, which has distinct interpretations. The knee parameter can be interpreted as the ‘timescale’ of the underlying data, whereby there is a direct mapping between the knee frequency as estimated from the aperiodic component and the decay time constant as estimated from the autocorrelation function (Miller et al., 2009; Gao et al., 2020). Based on this interpretation, N3 was found to have the lowest knee frequency, and as such would be interpreted as having the longest processing time, whereas REM sleep and wakefulness were found to have higher knee frequencies, and as such, faster processing timescales. These findings are consistent with other work that has examined aperiodic knees in sleep data (Lendner et al., 2023), autocorrelation analyses of sleep data (Zilio et al., 2021), as well as an analysis of timescales during sleep estimated from spiking activity (Hagemann et al., 2022).

Based on the pattern of findings and corresponding putative interpretations in this study, these results suggest that the general interpretation of sleep-related changes in aperiodic activity as a reflection of E-I balance may need to be amended to consider distinct patterns of changes across both E-I balance and neural timescales. Notably, while the pattern of E-I dynamics as inferred from the exponent between wake and sleep replicates, by fitting the knee model, a key finding is that, in intracranial data, it is the knee frequency that better discriminates between the different sleep stages, and relates to the changes in measured exponent when fitting a narrow / fixed model. This suggests that in addition to the global dynamics in E-I, a key change across sleep stages is changes in neural time scale – specifically that while EI dynamics may reflect the global dynamics between sleep and wake, shifts in timescale may additionally describe intra-sleep dynamics between sleep stages. This pattern is particularly clear in the intracranial data, in which measurements of neural timescale from local field potential (LFP) data have been established (Gao et al., 2020), but is also broadly consistent in the EEG data, suggesting that extracranial data also offers opportunities for measuring neural timescales.

Relatedly, by investigating sleeping data across both intra- and extracranial recordings, we highlight some notable commonalities as well as differences across modalities that should be further examined in future work. The ‘knee’ parameter, which has been much less studied in sleep research, was more prominently observed in the intracranial data, and less consistent in the extracranial EEG. This might relate to the relative scarcity of previous work focusing on aperiodic activity in sleep using iEEG, whereby the superior signal-to-noise ratio of intracranial data may be important for characterizing this feature of the data, allowing for further insights into the neural dynamics in sleep. However, it is also important to emphasize that despite the modality-related differences, the extracranial EEG data did show a clear knee in some stages, albeit less consistently. To our knowledge, this is one of the first clear demonstrations of aperiodic knees in extracranial data, and suggests ample opportunity for further investigating this aspect of the data, when employing analyses with frequency ranges and model forms that allow for capturing these features. Future work should further evaluate the occurrence and characteristics of ‘knees’ in sleep EEG data. This includes a thorough examination of how alterations in exponent measurements may correspond to underlying shifts in the knee across various sleep stages, as hinted at by the iEEG findings.

Another key aspect of this investigation is the use of time-resolved measures of aperiodic activity, extending beyond and complementing more common analyses of temporal patterns of oscillatory components (Stokes et al., 2023). In this study, for the first time to our knowledge, we showed that by tracking the temporal fluctuations of the spectral exponent, we are able to map the dynamics of transitions between distinct sleep stages, as well as track the sleeping brain’s event-related, transient, stimulus specific responses to auditory perturbations. This is consistent with contemporary work emphasizing the dynamic nature of aperiodic activity, with rapid changes across brain states and in response to external stimuli (Waschke et al, 2021; Wilson et al, 2022). Future work can further probe the temporal dynamics of aperiodic activity during sleep, which may offer potential for understanding the dynamic nature of neural activity during sleep and predicting spontaneous events (KCs, sleep spindles, slow waves, etc), as well as how how this activity relates to functional roles of aperiodic activity (Helfrich et al., 2021).

The findings here also have implications for sleep scoring – including suggesting features that may assist with detecting different sleep stages, and information about the transitions between sleep stages. While sleep scoring is defined primarily in terms of oscillatory features, this work supports previous work suggesting the aperiodic exponent may be a useful parameter for adjudicating between different sleep stages (Bodizs et al, 2021; Kozhemiako et al, 2022; Lendner et al, 2020). In addition, our findings extend previous results by showing that the aperiodic knee may also be especially useful in distinguishing between sleep stages, potentially contributing to improving the accuracy and reliability of the staging process. Particularly exciting is also the finding that time-resolved aperiodic parameters show distinct deviations over time during sleep-stage transitions, suggesting these measures may be useful to empirically investigate the progressions between sleep stages, and potentially offer increased time-resolution for detecting changes within and between sleep stages. Further research and validation across diverse sleep datasets should seek to examine the generalizability and reliability of these findings.

In analyzing neural responses to external events, we also observed transient, event-related steepening of the aperiodic exponent, particularly prominent during NREM sleep, which is consistent with an event-related increase in inhibition, possibly reflecting a response related the maintenance of sleep (Ameen et al., 2022; Blume et al., 2018). Notably, the exponent responses were also stimulus-specific, with an increased response following the presentation of unfamiliar voices as compared to familiar ones. These auditory-evoked exponent responses during sleep are consistent with our previous analyses showing a similar trend in brain responses during NREM sleep (Ameen et al., 2022). These similarities of the exponent responses to analyses of ERPs and elicited KCs raise the question of to what extent these features are independent and/or may reflect different measures of the same underlying events. Although our analyses suggest the presence of an aperiodic component contributing to the evoked KC, it’s important to note that the results are not conclusive. Changes in aperiodic exponent are still present when examining trials with no KCs, and are at least partially distinct from the ERP responses. These findings are consistent with other work showing event-related steepening of the exponent in response to auditory stimuli that is distinct from ERPs (Gyurkovics et al., 2022; Kałamała et al., 2023). Overall, while the analyses here do not fully establish the extent to which ERPs and exponent responses may reflect overlapping neural responses, these findings do establish event-related aperiodic responses during sleep that appear to be at least somewhat distinct to other common measures. Future work should further evaluate whether these inhibitory responses facilitate stimulus-specific processing and to what extent different evoked responses relate to each other.

## 5. Conclusion

In this investigation, we sought to explore aperiodic activity during sleep across both intra- and extracranial data. We showed that by using a broader frequency range and fitting a model with a knee, we can characterize more of the data and obtain estimates of aperiodic activity that map onto sleep stages, sleep transitions, and transient responses to external events. A key aspect of this work is the analysis of the knee frequency, which effectively distinguishes between sleep stages and offers potential new insights into the timescale of neuronal processing during sleep. Overall, these findings emphasize the relevance of examining aperiodic neural activity during sleep. By adopting these expanded parameters, we can gain fresh insights and perspectives on sleep processes, enhancing our understanding and interpretation of both the temporal and spectral dynamics of neural activity during sleep.

## Contributions

MSA, TD, JJ & KH: analysis plan. MS: experimental design and data collection. MSA and TD: data analysis. MSA and TD: manuscript drafting. All authors contributed to the manuscript.

## Acknowledgements

We would like to thank Christine Blume and Renate del Guidice for data collection. MSA is funded by the Austrian Academy of Sciences (ÖAW) and the Austrian Science Funds (FWF).

## 6. Supplements

### Supplemental Figures

**Figure 2-1.**
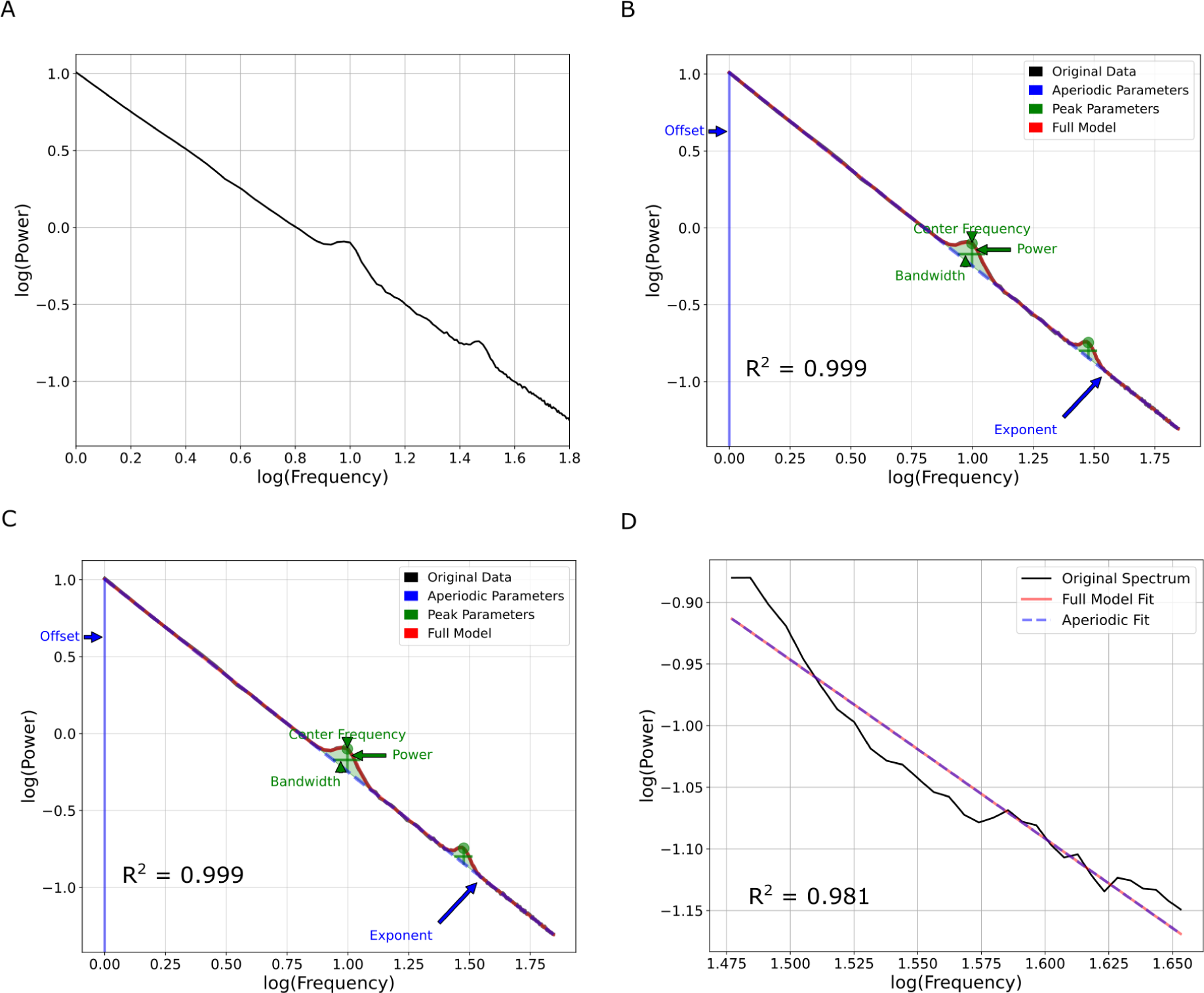
Simulated signal with no knee. A) Similar to Figure 1 we simulated a signal with two oscillatory peaks at 10 and 30 Hz. However, unlike Figure 1, we did not incorporate a knee. The knee model (B) and the fixed model with broad range (C) had comparable R^2^ and Exponent values. The fixed model with the narrow band (D) however, had lower R^2^ at 0.98. Note that even when no knee is present the knee model performed at high levels similar to the fixed model. R^2^: Goodness of fit.

**Figure 3-1.**
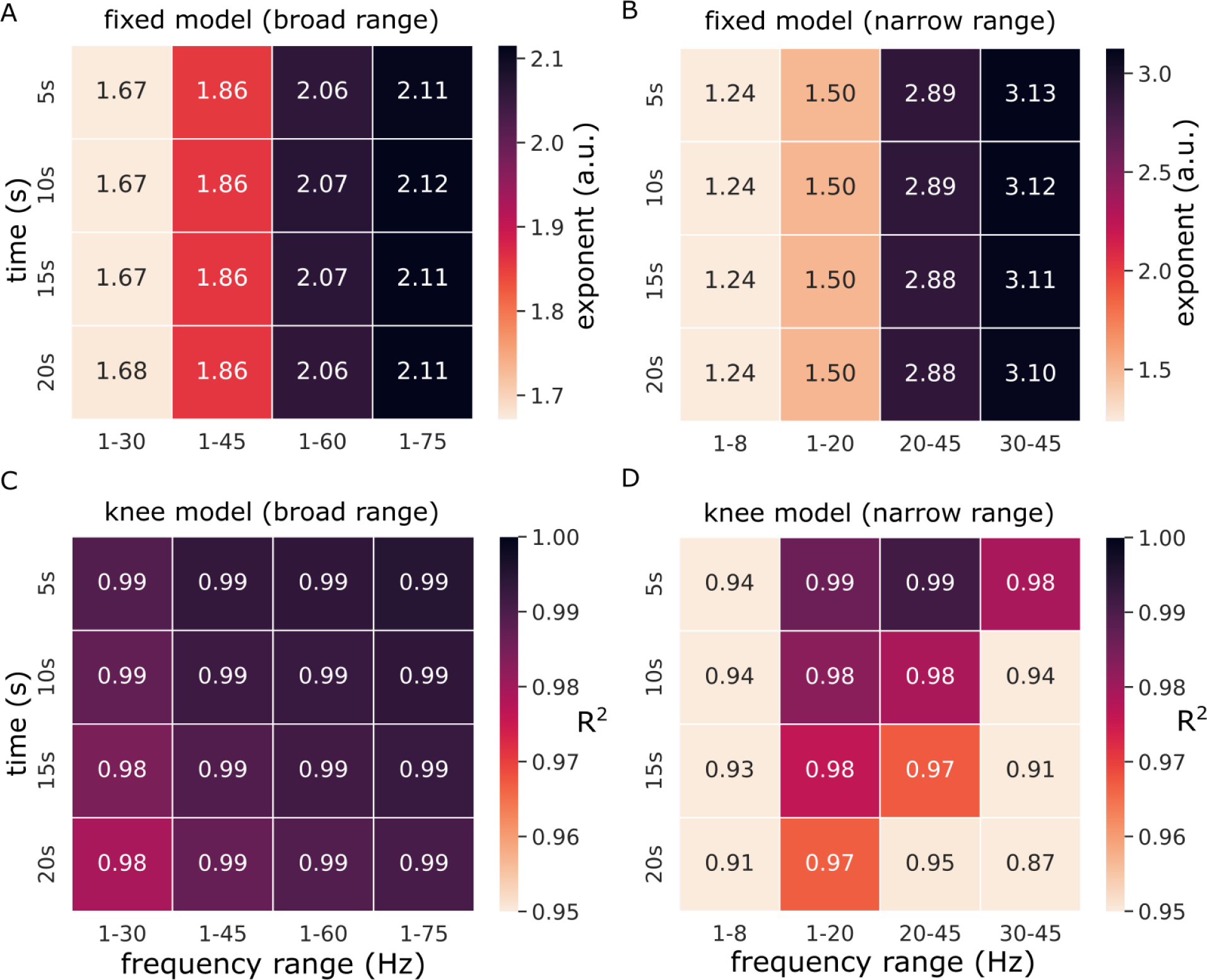
Model performance using broad and narrow frequency bands. A-B) The exponent values resulting from using A) broad frequency ranges and B) narrow frequency bands. Note the higher variance in (B) as compared to (A). C-D) The goodness of fit (R^2^) of the knee model fitted to the iEEG data using C) broadband or D) narrow frequency bands.

**Figure 3-2.**
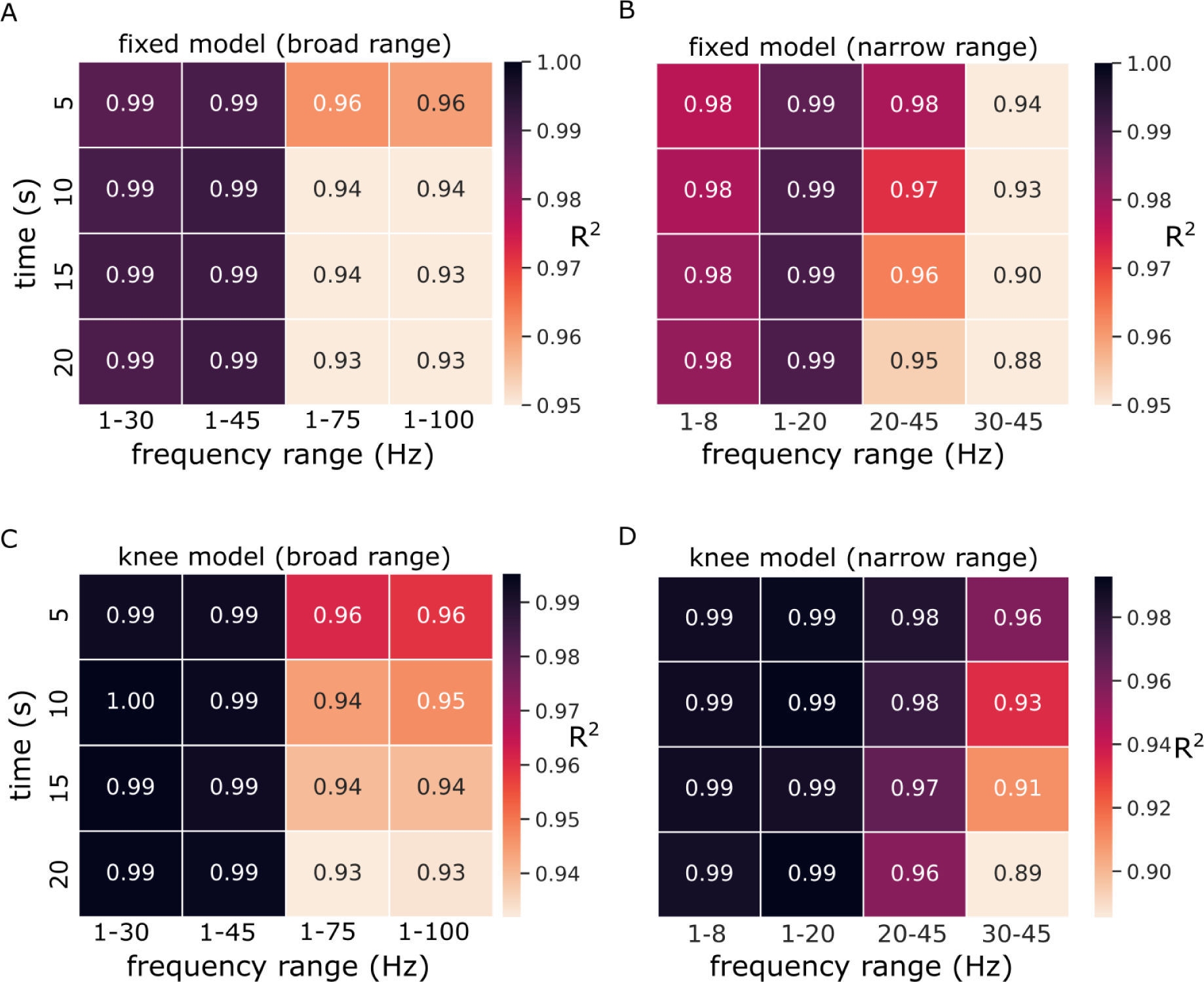
Model performance for different frequency ranges in EEG. A) The R^2^ values for the fixed model using broadband frequency ranges. B) The R^2^ values for the fixed model using narrow frequency ranges. C-D) The R^2^ values of the knee model using broadband (C) and narrowband (D) frequency ranges.

**Figure 4-1.**
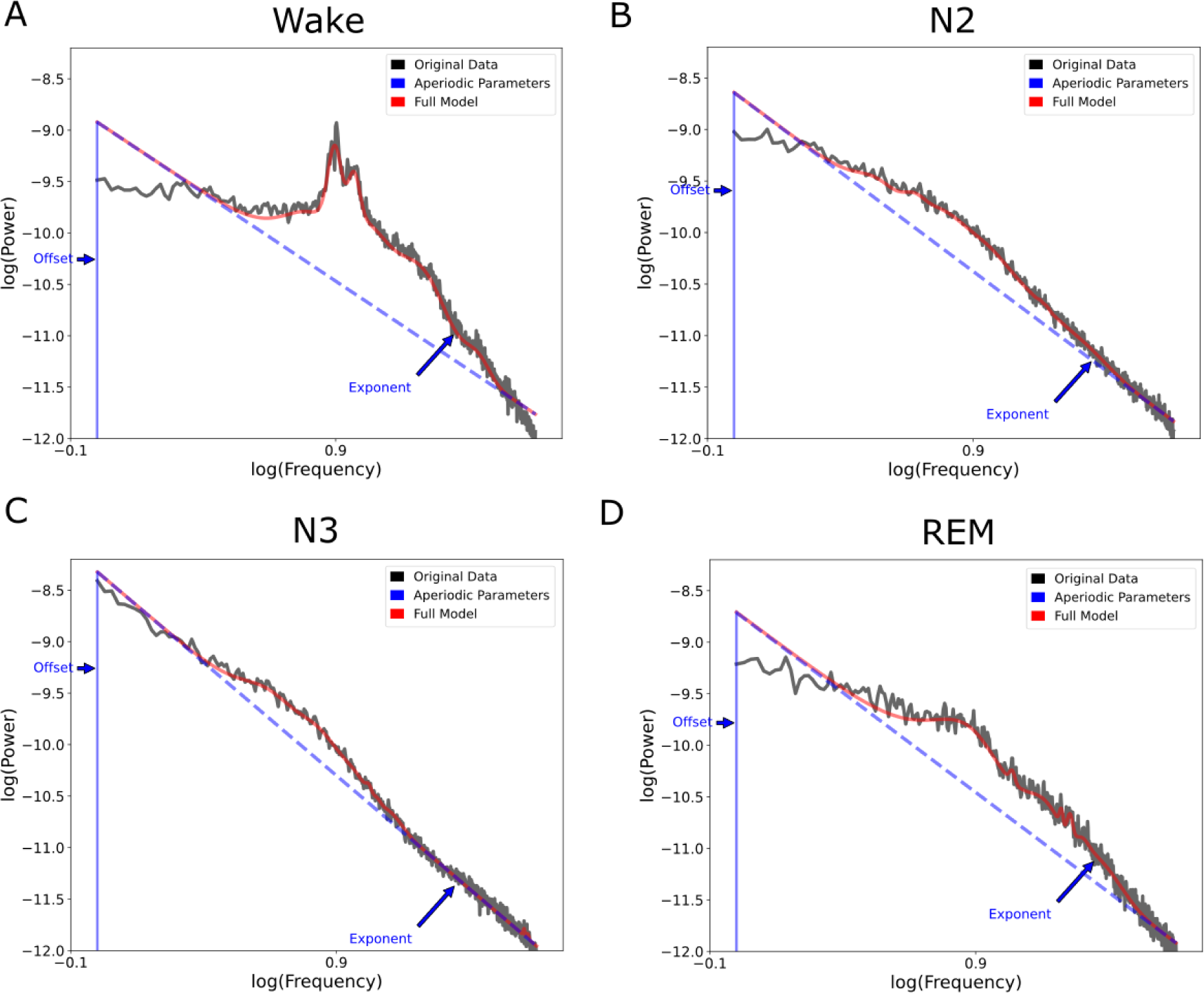
Model fit using a fixed model across a broad frequency range in the iEEG data. The fixed model is fitted to A) Wake, B) N2, C) N3, and D) REM sleep data.

**Figure 4-2.**
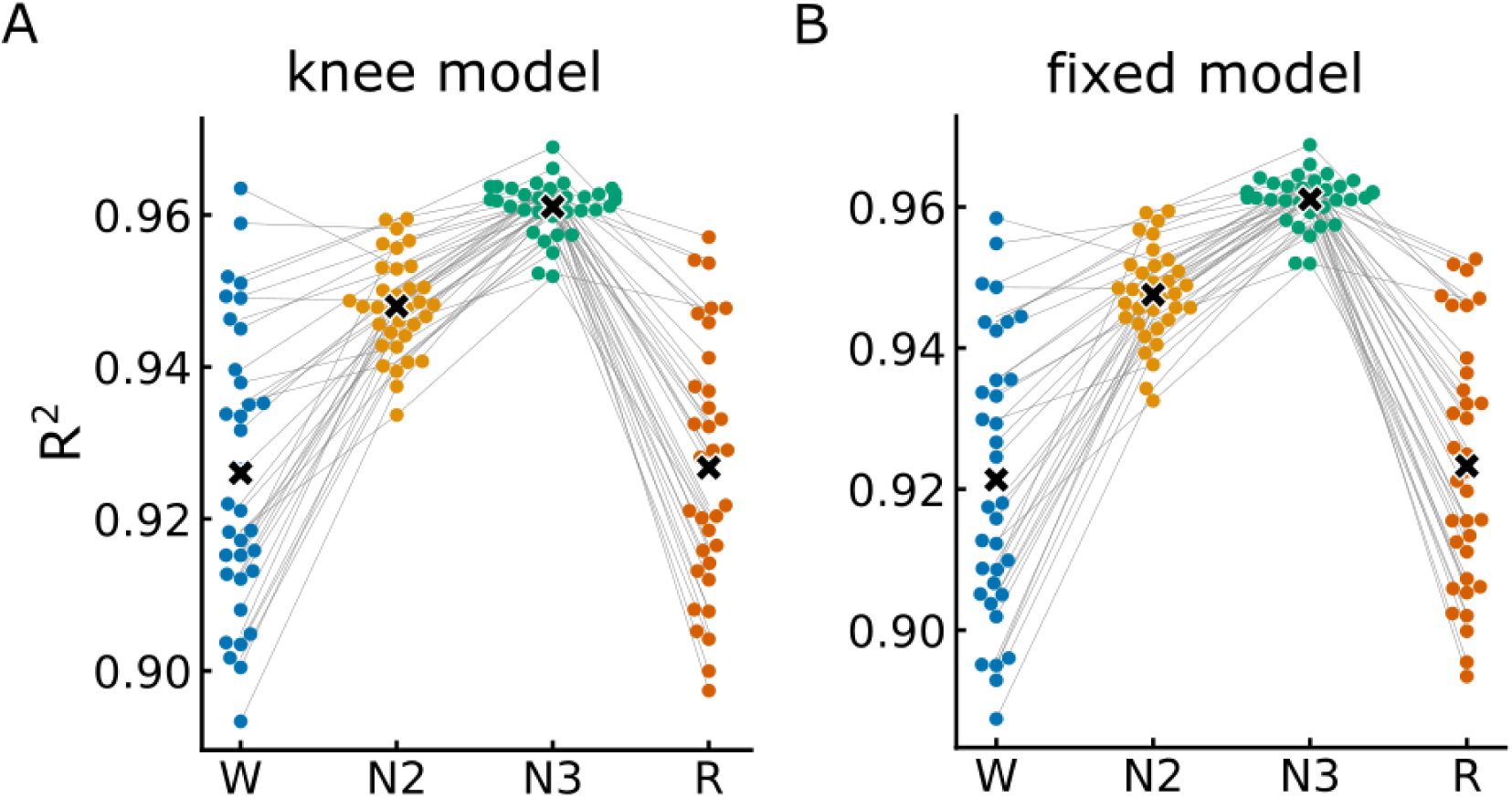
Model fit quality for the different sleep stages in the iEEG data. R^2^ of model fit using either A) the knee model (broad range) or B) fixed model (broad range). Note the overall high R^2^ values for all models and all stages (R^2^ > 0.86). Also note the high variability for Wake and REM stages as compared to N2 and N3.

**Figure 5-1.**
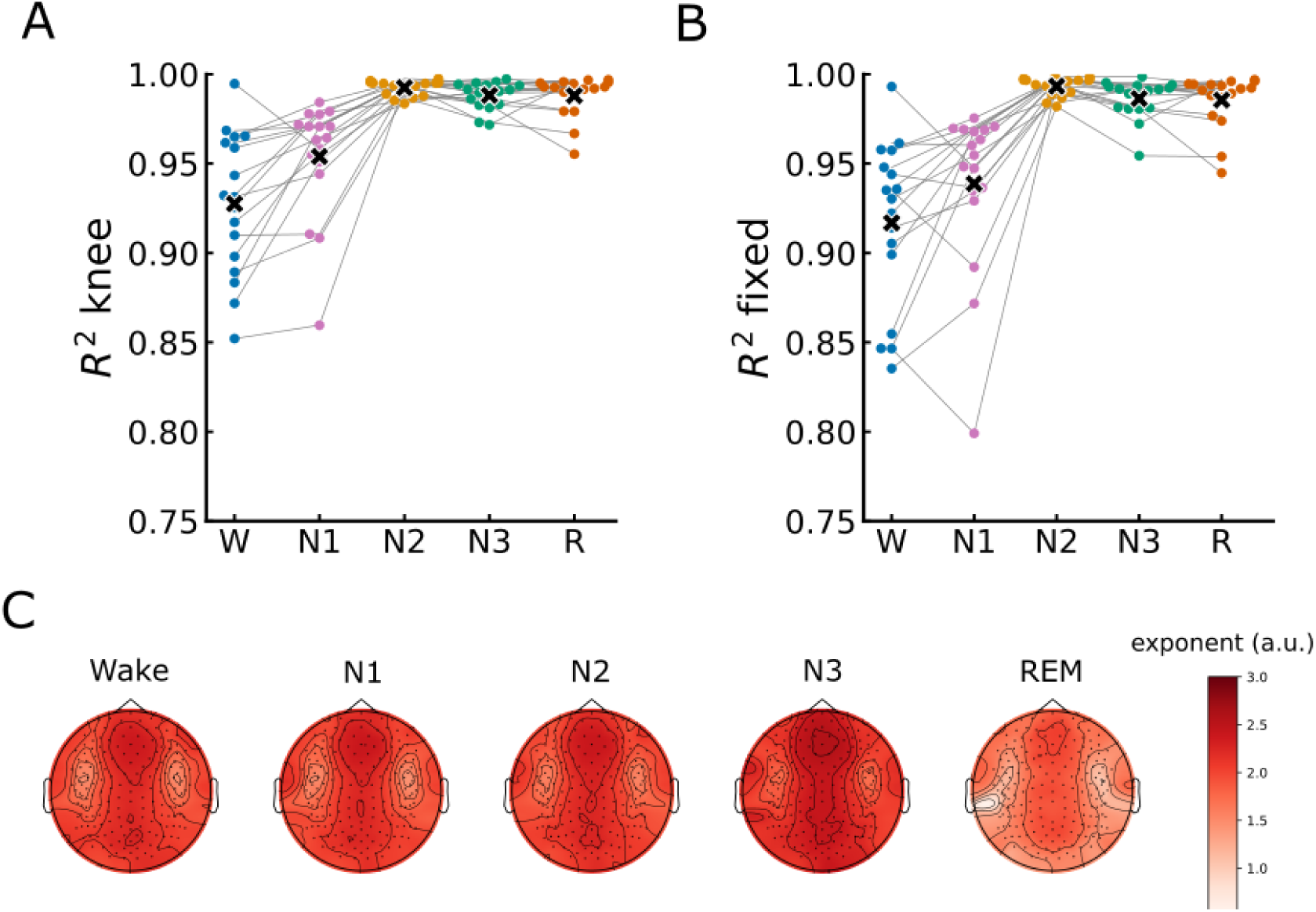
Extended EEG model fit results. A) R^2^ for the knee model for the different sleep stages. B) R^2^ of the fixed model for the different sleep stages. C) The topography of the exponent of the fixed model. Each dot in panels A and B represent one subject.

**Figure 6-1.**
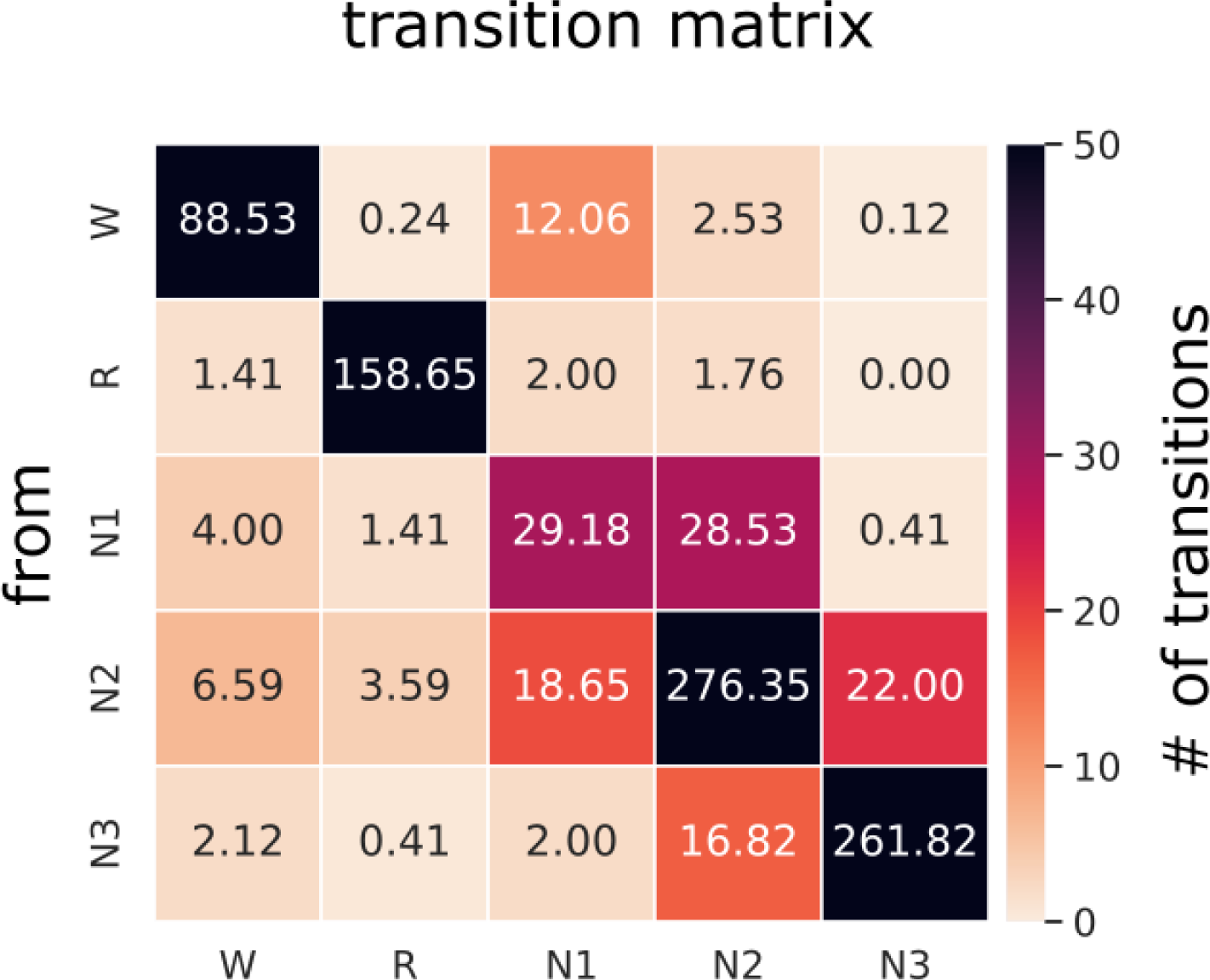
Matrix of transitions between sleep stages. Each cell represents the average number of transitions between two stages, averaged across all subjects.

**Figure 6-2.**
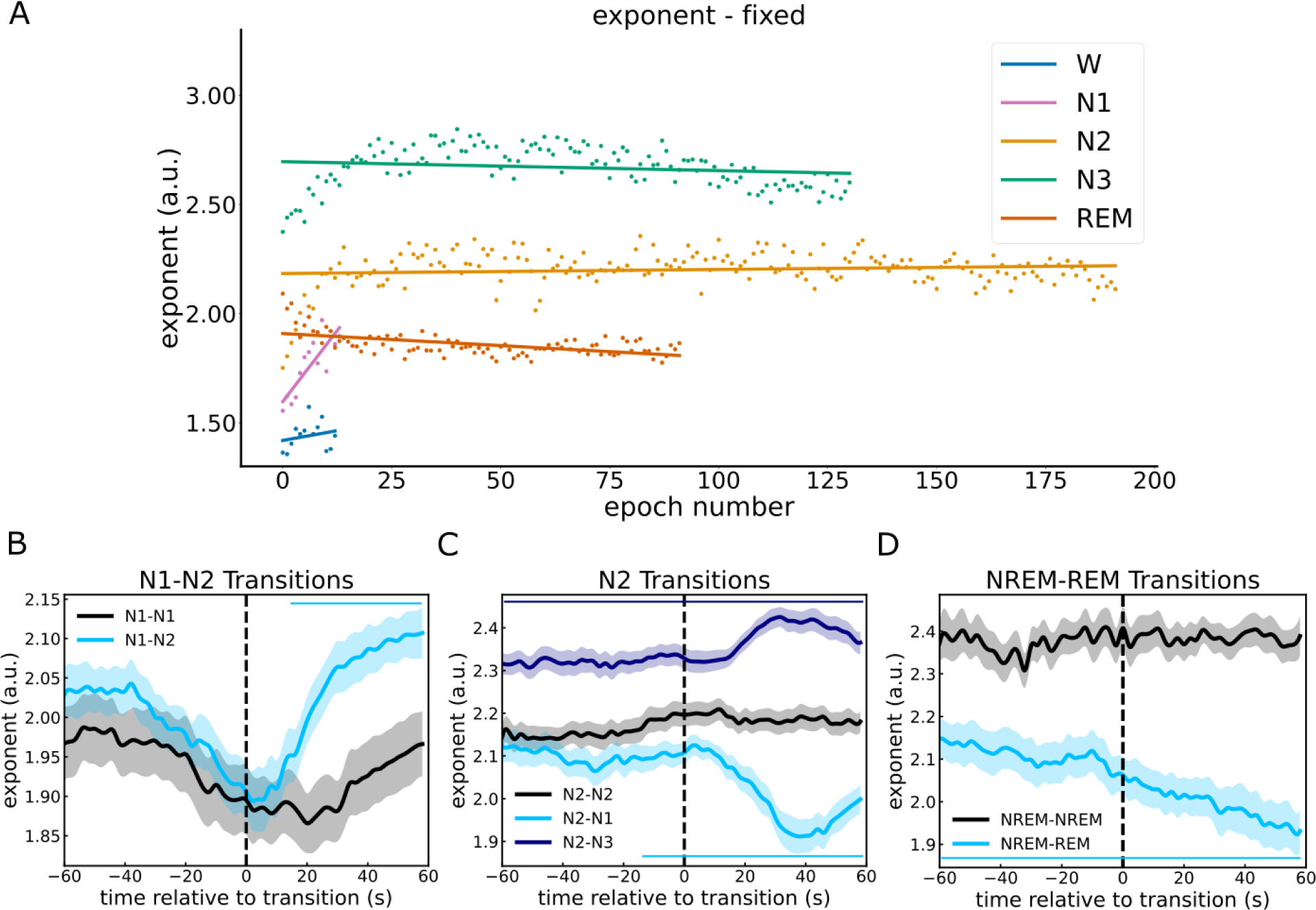
Replication of Figure 6 using the exponent of the fixed model. **A)** Epoch-by-epoch analysis of the exponent value throughout the night for each sleep stage. The results revealed for N1 and REM sleep the exponent changed significantly with the passage of time. The N1 exponent increased significantly N1: R^2^ = 0.7, F(1,12) = 27.59, p < 0.001) while that of REM decreased significantly (REM: R^2^ = 0.3, F(1,90) = 37.9, p = 0.01). For Wake, N2 and N3, however, the change in the exponent over time did not reach statistical significance (Wake: R^2^ = 0.05, F(1,11) = 0.54, p = 0.48 - N2: R^2^ = 0.01, F(1,190) = 0.54, p = 0.09 - N3: R^2^ = 0.03, F(1,29) = 3.58, p = 0.06). B-D) Exponent change around the transitions between sleep stages. B) Comparing the transitions from N1 to N2 against a baseline of continuous N1 showed that the exponent increased after the transition from N1 to N2 reaching a significant difference to that of the baseline starting at 14s following the transition (14s - 60s: ∑t(17) = 147.18, p < 0.001, d = 1.7). C) Examining the transitions from N2 to either N3 or N1. The results revealed a significant difference in the exponent between N2-N1 and N2-N2 that started 12s before the transition (-12s - 60s: ∑t(16) = 237.5, p < 0.001, d = -2.33). The exponent during N2 to N3 was significantly different from the baseline during the whole duration of the segment (-60s - 60s: ∑t(16) = 515.28, p < 0.001, d = 2.56), making it difficult to map the temporal dynamics of this transition. D) The exponent during the transitions from NREM to REM sleep showed a significant deviation from the baseline throughout the entire segment, with a substantial statistical difference (-60s - 60s: ∑t(16) = 335.16, p < 0.001, d = 1.69). These marked variances in C and D posed challenges in accurately mapping the temporal dynamics associated with this sleep stage transition.

**Figure 6-3.**
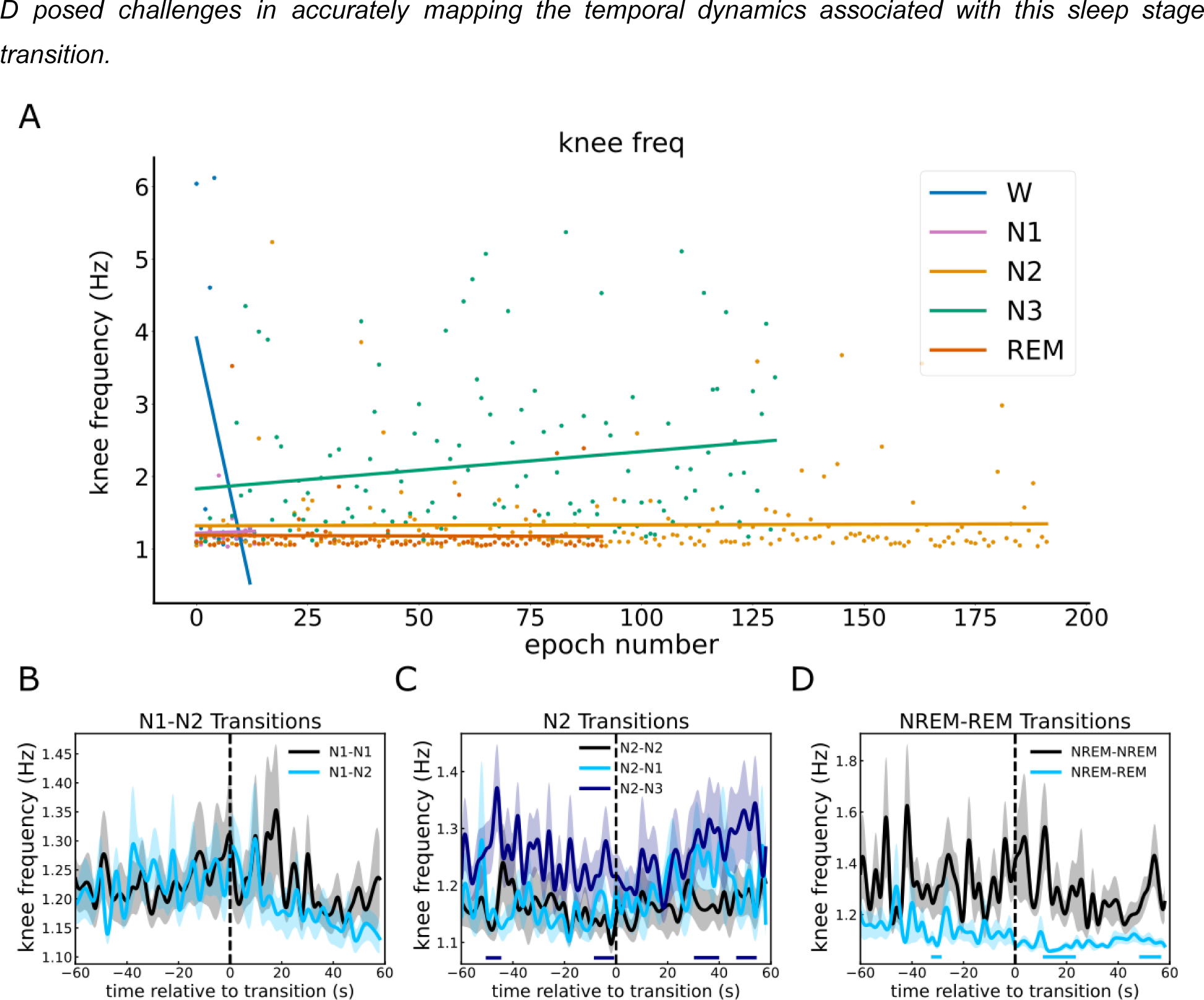
Replication of Figure 6 using the knee frequency. A) Epoch-by-epoch analysis of the exponent value throughout the night revealed a significant decrease in the knee frequency across wake epochs (R^2^ = 0.32, F(1,11) = 25.06, p = 0.046) and a significant increase across N3 epochs (R^2^ = 0.04, F(1,129) = 4.71, p = 0.03). N1, N2 and REM, however, showed no significant change in the knee frequency with the passage of time (N1: R^2^ = 0.001, F(1,12) = 0.01 p = 0.91, N2: R^2^ = 0.0002, F(1,190) = 27.59, p = 0.84, REM: R^2^ = 0.0002, F(1,90) = 0.02, = 0.89). B-D) Temporally resolved estimates of the knee frequency during transitions between sleep stages. B) No difference observed in the knee frequency during the transition from N1 to N2 as compared to a continuous N1 baseline (p > 0.1). C) The transitions from N2 to either N3 or N1. No difference between the transitions from N2 to N1 and the baseline (continuous N2, p > 0.05). For the transition from N2 to N3 however, four significant clusters were observed (-50s - -46s: ∑t(16) = 9.54, p = 0.01, d = 0.94, -6s - 0s: ∑t(16) = 9.98, p = 0.009, d = 0.78, 32s - 40s: ∑t(16) = 13.08, p = 0.004, d = 0.8, 48s - 54s: ∑t(16) = 11.42, p = 0.006, d = 0.88). D) NREM to REM transitions as compared to a NREM baseline differed significantly at three periods (-34s - 30s: ∑t(16) = 13.08, p = 0.004, d = 0.79, 12s - 22s: ∑t(16) = 15.18, p = 0.001, d = 0.88, 48s - 56s: ∑t(16) = 13.23, p = 0.004, d = 0.76)

**Figure 7-1.**
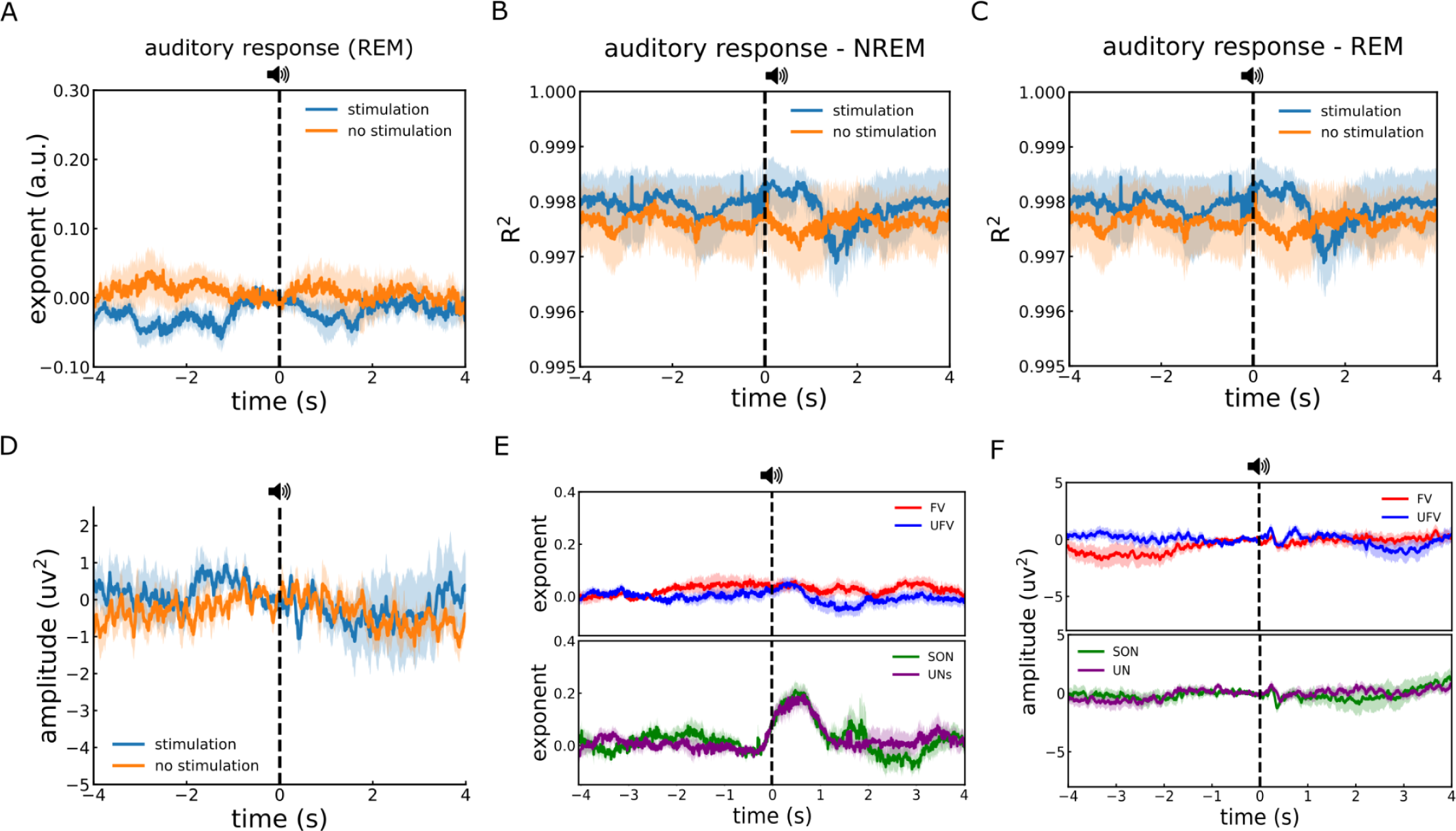
Auditory responses during REM sleep. A) Time-resolved estimates of aperiodic exponent in response to auditory stimuli, during REM sleep. B-C) The goodness of fits (R^2^) for fits around the presentation of auditory stimuli during (B) NREM and (C) REM. D-F) Auditory responses to different stimuli during REM sleep. D) Auditory ERPs during REM did not differ from a baseline of no stimulation. E-F) During REM sleep, when examining the responses to various voices and names, we found no significant differences in the exponent values for (E-top) different voices and (E-bottom) different names. Similarly, the ERPs also did not exhibit any notable variation between the different (F-top) voices and (F-bottom) names. Time zero represents stimulus onset. Dashed vertical line at time zero represents stimulus onset.

**Figure 7-2.**
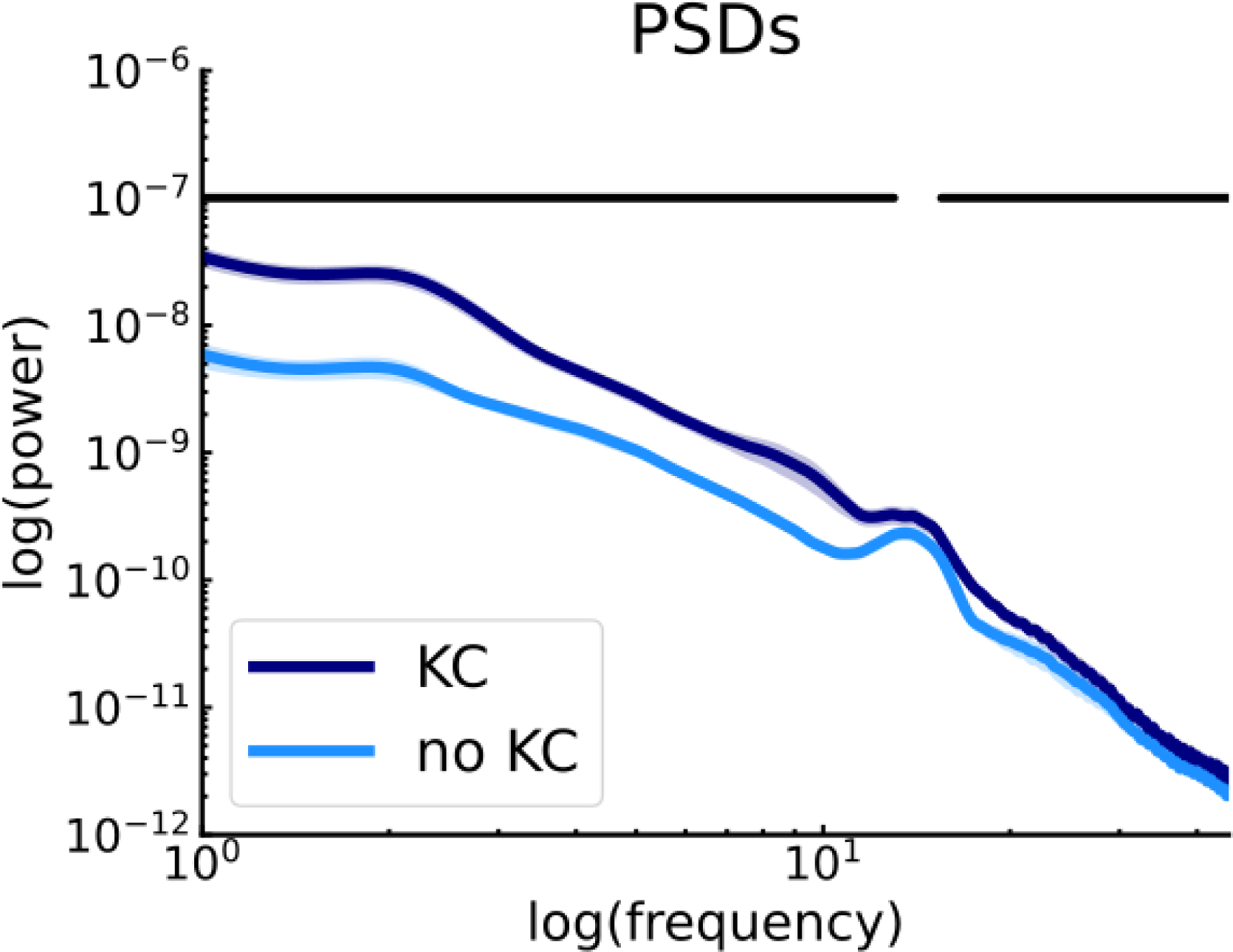
PSDs of the KC vs no KC analysis. We identified the peak points of exponent responses to auditory stimuli under two conditions: when a KC was evoked and when it was not. At these peak moments, we calculated the PSDs. It’s important to note the observed broadband variation in power, which was higher in trials where a KC was evoked. This suggests a shift in the aperiodic exponent rather than a difference that is attributed to oscillatory changes. The horizontal lines indicate the significant clusters.

## References

Alnes, S. L., Bächlin, L. Z. M., Schindler, K., & Tzovara, A. (2023). Neural complexity and the spectral slope characterise auditory processing in wakefulness and sleep. The European Journal of Neuroscience. 10.1111/ejn.16203

Ameen, M. S., Heib, D. P. J., Blume, C., & Schabus, M. (2022). The Brain Selectively Tunes to Unfamiliar Voices during Sleep. Journal of Neuroscience, 42(9), 1791–1803. 10.1523/JNEUROSCI.2524-20.2021

Ameen, M. S., Petzka, M., Peigneux, P., & Hoedlmoser, K. (2023). Post-training sleep modulates motor adaptation and task-related beta oscillations. Journal of Sleep Research, e14082. 10.1111/jsr.14082

American Association of Sleep Medicine, (AASM), & Iber, C. (2007). The AASM manual for the scoring of sleep and associated events: Rules, terminology, and specifications.

Anderer, P., Gruber, G., Parapatics, S., Woertz, M., Miazhynskaia, T., Klosch, G., Saletu, B., Zeitlhofer, J., Barbanoj, M. J., Danker-Hopfe, H., Himanen, S.-L., Kemp, B., Penzel, T., Grozinger, M., Kunz, D., Rappelsberger, P., Schlogl, A., & Dorffner, G. (2005). An E-health solution for automatic sleep classification according to Rechtschaffen and Kales: Validation study of the Somnolyzer 24 x 7 utilizing the Siesta database. Neuropsychobiology, 51(3), 115–133. 10.1159/000085205

Anderer, P., Moreau, A., Woertz, M., Ross, M., Gruber, G., Parapatics, S., Loretz, E., Heller, E., Schmidt, A., Boeck, M., Moser, D., Kloesch, G., Saletu, B., Saletu-Zyhlarz, G. M., Danker-Hopfe, H., Zeitlhofer, J., & Dorffner, G. (2010). Computer-assisted sleep classification according to the standard of the American Academy of Sleep Medicine: Validation study of the AASM version of the Somnolyzer 24 × 7. Neuropsychobiology, 62(4), 250–264. 10.1159/000320864

Berry, R. B., Brooks, R., Gamaldo, C., Harding, S. M., Lloyd, R. M., Quan, S. F., Troester, M. T., & Vaughn, B. V. (2017). AASM Scoring Manual Updates for 2017 (Version 2.4). Journal of Clinical Sleep Medicine: JCSM: Official Publication of the American Academy of Sleep Medicine, 13(5), 665–666. 10.5664/jcsm.6576

Bigdely-Shamlo, N., Mullen, T., Kothe, C., Su, K.-M., & Robbins, K. A. (2015). The PREP pipeline: Standardized preprocessing for large-scale EEG analysis. Frontiers in Neuroinformatics, 9, 16. 10.3389/fninf.2015.00016

Blume, C., Del Giudice, R., Wislowska, M., Heib, D. P. J., & Schabus, M. (2018). Standing sentinel during human sleep: Continued evaluation of environmental stimuli in the absence of consciousness. NeuroImage, 178, 638–648. 10.1016/j.neuroimage.2018.05.056

Bódizs, R., Szalárdy, O., Horváth, C., Ujma, P. P., Gombos, F., Simor, P., Pótári, A., Zeising, M., Steiger, A., & Dresler, M. (2021). A set of composite, non-redundant EEG measures of NREM sleep based on the power law scaling of the Fourier spectrum. Scientific Reports, 11(1). 10.1038/s41598-021-81230-7

Bridi, M. C. D., Zong, F.-J., Min, X., Luo, N., Tran, T., Qiu, J., Severin, D., Zhang, X.-T., Wang, G., Zhu, Z.-J., He, K.-W., & Kirkwood, A. (2020). Daily Oscillation of the Excitation-Inhibition Balance in Visual Cortical Circuits. Neuron, 105(4), 621–629.e4. 10.1016/j.neuron.2019.11.011

Chini, M., Pfeffer, T., & Hanganu-Opatz, I. (2022). An increase of inhibition drives the developmental decorrelation of neural activity. eLife, 11, e78811. 10.7554/eLife.78811

Delorme, A., & Makeig, S. (2004). EEGLAB: An open source toolbox for analysis of single-trial EEG dynamics including independent component analysis. Journal of Neuroscience Methods, 134(1), 9–21. 10.1016/j.jneumeth.2003.10.009

Demirel, B. U., Skelin, I., Zhang, H., Lin, J. J., & Abdullah Al Faruque, M. (2021). Single-Channel EEG Based Arousal Level Estimation Using Multitaper Spectrum Estimation at Low-Power Wearable Devices. 2021 43rd Annual International Conference of the IEEE Engineering in Medicine & Biology Society (EMBC), 542–545. 10.1109/EMBC46164.2021.9629733

Donoghue, T. (2019). LISC: A Python Package for Scientific Literature Collection and Analysis. Journal of Open Source Software, 4(41), 1674. 10.21105/joss.01674

Donoghue, T., Haller, M., Peterson, E. J., Varma, P., Sebastian, P., Gao, R., Noto, T., Lara, A. H., Wallis, J. D., Knight, R. T., Shestyuk, A., & Voytek, B. (2020). Parameterizing neural power spectra into periodic and aperiodic components. Nature Neuroscience, 23(12), 1655–1665. 10.1038/s41593-020-00744-x

Donoghue, T., & Watrous, A. J. (2023). How Can We Differentiate Narrow-Band Oscillations from Aperiodic Activity? In N. Axmacher (Ed.), Intracranial EEG: A Guide for Cognitive Neuroscientists (pp. 351–364). Springer International Publishing. 10.1007/978-3-031-20910-9_22

Favaro, J., Colombo, M. A., Mikulan, E., Sartori, S., Nosadini, M., Pelizza, M. F., Rosanova, M., Sarasso, S., Massimini, M., & Toldo, I. (2023). The maturation of aperiodic EEG activity across development reveals a progressive differentiation of wakefulness from sleep. NeuroImage, 277, 120264. 10.1016/j.neuroimage.2023.120264

Feinberg, I., March, J. D., Floyd, T. C., Fein, G., & Aminoff, M. J. (1984). Log amplitude is a linear function of log frequency in NREM sleep EEG of young and elderly normal subjects. Electroencephalography and Clinical Neurophysiology, 58(2), 158–160. 10.1016/0013-4694(84)90029-4

Foygel, R., & Drton, M. (2010). Extended Bayesian information criteria for Gaussian graphical models. Proceedings of the 23rd International Conference on Neural Information Processing Systems - Volume 1, 604–612.

Frauscher, B., von Ellenrieder, N., Zelmann, R., Doležalová, I., Minotti, L., Olivier, A., Hall, J., Hoffmann, D., Nguyen, D. K., Kahane, P., Dubeau, F., & Gotman, J. (2018). Atlas of the normal intracranial electroencephalogram: Neurophysiological awake activity in different cortical areas. Brain, 141(4), 1130–1144. 10.1093/brain/awy035

Freeman, W. J., & Zhai, J. (2009). Simulated power spectral density (PSD) of background electrocorticogram (ECoG). Cognitive Neurodynamics, 3(1), 97–103. 10.1007/s11571-008-9064-y

G. Horváth, C., Szalárdy, O., Ujma, P. P., Simor, P., Gombos, F., Kovács, I., Dresler, M., & Bódizs, R. (2022). Overnight dynamics in scale-free and oscillatory spectral parameters of NREM sleep EEG. Scientific Reports, 12(1). 10.1038/s41598-022-23033-y

Gao, R., Peterson, E. J., & Voytek, B. (2017). Inferring synaptic excitation/inhibition balance from field potentials. NeuroImage, 158, 70–78. 10.1016/j.neuroimage.2017.06.078

Gao, R., van den Brink, R. L., Pfeffer, T., & Voytek, B. (2020). Neuronal timescales are functionally dynamic and shaped by cortical microarchitecture. eLife, 9, e61277. 10.7554/eLife.61277

Gerster, M., Waterstraat, G., Litvak, V., Lehnertz, K., Schnitzler, A., Florin, E., Curio, G., & Nikulin, V. (2022). Separating Neural Oscillations from Aperiodic 1/f Activity: Challenges and Recommendations. Neuroinformatics, 20(4), 991–1012. 10.1007/s12021-022-09581-8

Gramfort, A., Luessi, M., Larson, E., Engemann, D., Strohmeier, D., Brodbeck, C., Goj, R., Jas, M., Brooks, T., Parkkonen, L., & Hämäläinen, M. (2013). MEG and EEG data analysis with MNE-Python. Frontiers in Neuroscience, 7. https://www.frontiersin.org/articles/10.3389/fnins.2013.00267

Gyurkovics, M., Clements, G. M., Low, K. A., Fabiani, M., & Gratton, G. (2022). Stimulus-Induced Changes in 1/f-like Background Activity in EEG. Journal of Neuroscience, 42(37), 7144–7151. 10.1523/JNEUROSCI.0414-22.2022

Hagemann, A., Kehl, M. S., Dehning, J., Spitzner, F. P., Niediek, J., Wibral, M., Mormann, F., & Priesemann, V. (2022). Intrinsic timescales of spiking activity in humans during wakefulness and sleep (arXiv:2205.10308). arXiv. http://arxiv.org/abs/2205.10308

He, B. J. (2014). Scale-free brain activity: Past, present, and future. Trends in Cognitive Sciences, 18(9), 480–487. 10.1016/j.tics.2014.04.003

He, B. J., Zempel, J. M., Snyder, A. Z., & Raichle, M. E. (2010). The temporal structures and functional significance of scale-free brain activity. Neuron, 66(3), 353–369. 10.1016/j.neuron.2010.04.020

Helfrich, R. F., Lendner, J. D., & Knight, R. T. (2021). Aperiodic sleep networks promote memory consolidation. Trends in Cognitive Sciences, 25(8), 648–659. 10.1016/j.tics.2021.04.009

Höhn, C., Hahn, M. A., Lendner, J. D., & Hoedlmoser, K. (2023). Spectral slope and Lempel-Ziv complexity as robust markers of brain states during sleep and wakefulness (p. 2022.09.10.507390). bioRxiv. 10.1101/2022.09.10.507390

Kałamała, P., Gyurkovics, M., Bowie, D. C., Clements, G. M., Low, K. A., Dolcos, F., Fabiani, M., & Gratton, G. (2023). Event-Induced Modulation of Aperiodic Background EEG: Attention-Dependent and Age-Related Shifts in E:I balance, and Their Consequences for Behavior (p. 2023.04.14.536436). bioRxiv. 10.1101/2023.04.14.536436

Kozhemiako, N., Mylonas, D., Pan, J. Q., Prerau, M. J., Redline, S., & Purcell, S. M. (2022). Sources of Variation in the Spectral Slope of the Sleep EEG. eNeuro, 9(5), ENEURO.0094-22.2022. 10.1523/ENEURO.0094-22.2022

Lendner, J. D., Helfrich, R. F., Mander, B. A., Romundstad, L., Lin, J. J., Walker, M. P., Larsson, P. G., & Knight, R. T. (2020). An electrophysiological marker of arousal level in humans. eLife, 9, e55092. 10.7554/eLife.55092

Lendner, J. D., Niethard, N., Mander, B. A., van Schalkwijk, F. J., Schuh-Hofer, S., Schmidt, H., Knight, R. T., Born, J., Walker, M. P., Lin, J. J., & Helfrich, R. F. (2023). Human REM sleep recalibrates neural activity in support of memory formation. Science Advances, 9(34), eadj1895. 10.1126/sciadv.adj1895

Lombardi, F., Herrmann, H. J., & de Arcangelis, L. (2017). Balance of excitation and inhibition determines 1/f power spectrum in neuronal networks. Chaos (Woodbury, N.Y.), 27(4), 047402. 10.1063/1.4979043

Maris, E., & Oostenveld, R. (2007). Nonparametric statistical testing of EEG- and MEG-data. Journal of Neuroscience Methods, 164(1), 177–190. 10.1016/j.jneumeth.2007.03.024

Maschke, C., Duclos, C., Owen, A. M., Jerbi, K., & Blain-Moraes, S. (2023). Aperiodic brain activity and response to anesthesia vary in disorders of consciousness. NeuroImage, 275, 120154. 10.1016/j.neuroimage.2023.120154

Miller, K. J., Sorensen, L. B., Ojemann, J. G., & Nijs, M. den. (2009). Power-Law Scaling in the Brain Surface Electric Potential. PLOS Computational Biology, 5(12), e1000609. 10.1371/journal.pcbi.1000609

Miskovic, V., MacDonald, K. J., Rhodes, L. J., & Cote, K. A. (2019). Changes in EEG multiscale entropy and power-law frequency scaling during the human sleep cycle. Human Brain Mapping, 40(2), 538–551. 10.1002/hbm.24393

Niethard, N., Hasegawa, M., Itokazu, T., Oyanedel, C. N., Born, J., & Sato, T. R. (2016). Sleep-Stage-Specific Regulation of Cortical Excitation and Inhibition. Current Biology: CB, 26(20), 2739–2749. 10.1016/j.cub.2016.08.035

Oostenveld, R., Fries, P., Maris, E., & Schoffelen, J.-M. (2010). FieldTrip: Open Source Software for Advanced Analysis of MEG, EEG, and Invasive Electrophysiological Data. Computational Intelligence and Neuroscience, 2011, 156869. 10.1155/2011/156869

Parapatics, S., Anderer, P., Gruber, G., Saletu, B., Saletu-Zyhlarz, G., & Dorffner, G. (2015). K-complex amplitude as a marker of sleep homeostasis in obstructive sleep apnea syndrome and healthy controls. Somnologie - Schlafforschung Und Schlafmedizin, 19(1), 22–29. 10.1007/s11818-015-0701-5

Pereda, E., Gamundi, A., Rial, R., & González, J. (1998). Non-linear behaviour of human EEG: Fractal exponent versus correlation dimension in awake and sleep stages. Neuroscience Letters, 250(2), 91–94. 10.1016/S0304-3940(98)00435-2

Raftery, A. E. (1995). Bayesian Model Selection in Social Research. Sociological Methodology, 25, 111–163. 10.2307/271063

Rosenblum, Y., Bovy, L., Weber, F. D., Steiger, A., Zeising, M., & Dresler, M. (2022). Increased Aperiodic Neural Activity During Sleep in Major Depressive Disorder. Biological Psychiatry Global Open Science. 10.1016/j.bpsgos.2022.10.001

Schneider, B., Szalárdy, O., Ujma, P. P., Simor, P., Gombos, F., Kovács, I., Dresler, M., & Bódizs, R. (2022). Scale-free and oscillatory spectral measures of sleep stages in humans. Frontiers in Neuroinformatics, 16, 989262. 10.3389/fninf.2022.989262

Schwarz, J. F. A., Åkerstedt, T., Lindberg, E., Gruber, G., Fischer, H., & Theorell-Haglöw, J. (2017). Age affects sleep microstructure more than sleep macrostructure. Journal of Sleep Research, 26(3), 277–287. 10.1111/jsr.12478

Seabold, S., & Perktold, J. (2010). Statsmodels: Econometric and Statistical Modeling with Python. 92–96. 10.25080/Majora-92bf1922-011

Stokes, P. A., Rath, P., Possidente, T., He, M., Purcell, S., Manoach, D. S., Stickgold, R., & Prerau, M. J. (2023). Transient oscillation dynamics during sleep provide a robust basis for electroencephalographic phenotyping and biomarker identification. Sleep, 46(1), zsac223. 10.1093/sleep/zsac223

Trakoshis, S., Martínez-Cañada, P., Rocchi, F., Canella, C., You, W., Chakrabarti, B., Ruigrok, A. N., Bullmore, E. T., Suckling, J., Markicevic, M., Zerbi, V., MRC AIMS Consortium, Baron-Cohen, S., Gozzi, A., Lai, M.-C., Panzeri, S., & Lombardo, M. V. (2020). Intrinsic excitation-inhibition imbalance affects medial prefrontal cortex differently in autistic men versus women. eLife, 9, e55684. 10.7554/eLife.55684

von Ellenrieder, N., Gotman, J., Zelmann, R., Rogers, C., Nguyen, D. K., Kahane, P., Dubeau, F., & Frauscher, B. (2020). How the Human Brain Sleeps: Direct Cortical Recordings of Normal Brain Activity. Annals of Neurology, 87(2), 289–301. 10.1002/ana.25651

Voytek, B., Kramer, M. A., Case, J., Lepage, K. Q., Tempesta, Z. R., Knight, R. T., & Gazzaley, A. (2015). Age-Related Changes in 1/f Neural Electrophysiological Noise. Journal of Neuroscience, 35(38), 13257–13265. 10.1523/JNEUROSCI.2332-14.2015

Waschke, L., Donoghue, T., Fiedler, L., Smith, S., Garrett, D. D., Voytek, B., & Obleser, J. (2021). Modality-specific tracking of attention and sensory statistics in the human electrophysiological spectral exponent. eLife, 10, e70068. 10.7554/eLife.70068

Welch, P. (1967). The use of fast Fourier transform for the estimation of power spectra: A method based on time averaging over short, modified periodograms. IEEE Transactions on Audio and Electroacoustics, 15(2), 70–73. 10.1109/TAU.1967.1161901

Wen, H., & Liu, Z. (2016). Separating Fractal and Oscillatory Components in the Power Spectrum of Neurophysiological Signal. Brain Topography, 29(1), 13–26. 10.1007/s10548-015-0448-0

Wilson, L. E., da Silva Castanheira, J., & Baillet, S. (2022). Time-resolved parameterization of aperiodic and periodic brain activity. eLife, 11, e77348. 10.7554/eLife.77348

Woertz, M., Miazhynskaia, T., Anderer, P., & Dorffner, G. (2004). Automatic K-complex detection: Comparison of two different approaches. Abstracts of the ESRS, JSR 13(Suppl.1),1, Prague 2004.

Zilio, F., Gomez-Pilar, J., Cao, S., Zhang, J., Zang, D., Qi, Z., Tan, J., Hiromi, T., Wu, X., Fogel, S., Huang, Z., Hohmann, M. R., Fomina, T., Synofzik, M., Grosse-Wentrup, M., Owen, A. M., & Northoff, G. (2021). Are intrinsic neural timescales related to sensory processing? Evidence from abnormal behavioral states. NeuroImage, 226, 117579. 10.1016/j.neuroimage.2020.117579

